# BCLAF1 links RNA splicing to ATF4-dependent metabolic adaptation in acute myeloid leukemia

**DOI:** 10.64898/2026.01.19.700325

**Authors:** Laura López-Hernández, Stephanie J. Crowley, Sara Cea-Sánchez, Ricardo de Arellano, Neetij Krishnan, Patrick Toolan-Kerr, Lynn White, Domenico Ignoti, Emily Soto-Hidalgo, Stefano Gustincich, Román González-Prieto, Luca Pandolfini, Isaia Barbieri, Patricia Altea-Manzano, Jeffrey A. Magee, Jeffrey J Bednarski, Gonzalo Millán-Zambrano

**Author notes:** Correspondence: Jeffrey J. Bednarski; Washington University School of Medicine; Department of Pediatrics, 660 S. Euclid Ave., Campus Box 8208, St. Louis, MO 63110, USA., Gonzalo Millán-Zambrano; Centro Andaluz de Biología Molecular y Medicina Regenerativa-CABIMER, CSIC-Universidad de Sevilla-Universidad Pablo de Olavide, 41092 Seville, Spain. These authors contributed equally (Laura López-Hernández, Stephanie J. Crowley).

## Abstract

Acute myeloid leukemia (AML) is driven by a combination of genetic alterations and non-mutational mechanisms that disrupt normal hematopoiesis and support leukemic cell survival. While the mutational landscape of AML is well characterized, the non-genetic processes that sustain leukemic maintenance remain comparatively less understood. Using human AML cell lines and murine models of AML, we identify BCL2-associated transcription factor 1 (BCLAF1) as a key regulator of leukemic progression through control of mRNA processing. BCLAF1 physically associates with core spliceosome components and regulates alternative splicing, with a predominant effect on intron retention. We demonstrate that BCLAF1 is required for the productive splicing of activating transcription factor 4 (ATF4) mRNA, thereby sustaining ATF4 protein expression. Loss of BCLAF1 reduces ATF4 protein levels, leading to downregulation of metabolic target genes and disruption of *de novo* amino acid biosynthesis. Furthermore, depletion of BCLAF1 sensitizes AML cells to venetoclax, a clinically relevant BCL-2 inhibitor. Together, these findings uncover a previously unrecognized role for BCLAF1 in coordinating mRNA splicing and metabolic adaptation in AML, highlighting its potential as a therapeutic target.

**Statement of significance:** Aberrant RNA splicing and metabolic reprogramming are hallmarks of cancer, yet how these processes are mechanistically linked remains unclear. This study identifies BCLAF1 as a key regulator connecting splicing control to amino acid metabolism in acute myeloid leukemia, revealing a previously unrecognized functional vulnerability at the intersection of these pathways.

## Introduction

Acute myeloid leukemia (AML) is an aggressive hematologic malignancy characterized by the clonal expansion of immature myeloid precursor cells. While many patients initially respond to chemotherapy, most eventually relapse due to the persistence of drug-resistant leukemic stem cells (LSCs). (1,2) Despite recent therapeutic advances, AML remains largely incurable, with a 5-year overall survival rate of about 20%,(3) underscoring the need for deeper mechanistic insights and new therapeutic targets. (4)

AML is a biologically heterogeneous disease driven by a wide array of somatic oncogenic mutations. Among these, mutations in spliceosome genes, present in 10-25% of cases, are particularly prominent.(5,6) Although their precise oncogenic roles remain incompletely understood, growing evidence implicates aberrant mRNA splicing as a critical mediator of leukemogenesis.(7,8) Notably, AML cells bearing spliceosome mutations exhibit selective sensitivity to perturbations of the splicing machinery,(9–12) highlighting the spliceosome as a promising therapeutic target.

By contrast, the non-mutational molecular processes that sustain AML are less well defined. One such process is metabolic reprogramming, which has recently emerged as a hallmark of leukemogenesis. (13,14) LSCs exhibit elevated amino acid uptake and catabolism to support oxidative phosphorylation.(15) In addition, AML cells display specific dependencies on distinct amino acids,(16–21) underscoring the therapeutic potential of targeting amino acid metabolism. However, the upstream regulatory mechanisms that coordinate these metabolic adaptations remain poorly defined.

Emerging evidence indicates that mRNA splicing and cellular metabolism are interconnected. Metabolic cues extensively regulate mRNA splicing programs, enabling rapid cellular adaptation to nutrient availability and stress. (22) Conversely, recent work suggests that perturbation of the splicing machinery can reshape cellular metabolic states, particularly in cancer cells. (23,24) However, whether splicing dysregulation directly enforces metabolic reprogramming required for leukemic cell survival remains largely unexplored.

Here, we uncover a central role for BCLAF1 in coupling RNA processing with metabolic programs in AML. We show that BCLAF1 promotes productive mRNA splicing in AML cells and identify the metabolic regulator ATF4 as a key downstream effector. Loss of BCLAF1 disrupts *ATF4* mRNA processing, leading to reduced ATF4 protein levels and down-regulation of amino acid metabolic genes. These findings establish BCLAF1 as a crucial regulator of mRNA processing in AML and reveal a mechanistic link between splicing control and metabolic adaptation supporting leukemic cell survival.

## Results

### BCLAF1 supports AML pathogenesis

Cancer dependencies can be predicted based on gene expression, (25) offering a promising strategy for identifying clinically relevant biomarkers and therapeutic targets. Based on our previous work identifying BCLAF1 as a context-dependent regulator of hematopoietic progenitor–associated transcriptional programs, (26) we explored whether BCLAF1 might also contribute to leukemic cell states in AML. To assess the clinical relevance of BCLAF1 in human cancer, we analyzed publicly available gene expression data from The Cancer Genome Atlas (TCGA). BCLAF1 is most highly expressed in AML samples compared to other cancer types (Figure 1A) and is significantly elevated in AML relative to healthy hematopoietic tissues (Figure 1B), suggesting that BCLAF1 may play a central role in AML. To directly assess this possibility, we performed growth competition assays in two independent human *MLL-AF9* rearranged AML cell lines, MOLM-13 (*FLT3-ITD*) and THP-1 (*FLT3* wild-type). Cells were transduced with constructs co-expressing GFP, Cas9, and either a non-targeting negative control sgRNA, one of two independent *BCLAF1*-targeting sgRNAs, or a positive control sgRNA against *RBM39*, which is essential for AML maintenance (27). In both cell lines, we observed a substantial depletion of GFP-positive cells expressing *BCLAF1* sgRNAs (Figure 1C), indicating that BCLAF1 is required for human AML cell proliferation.

**Figure 1.**
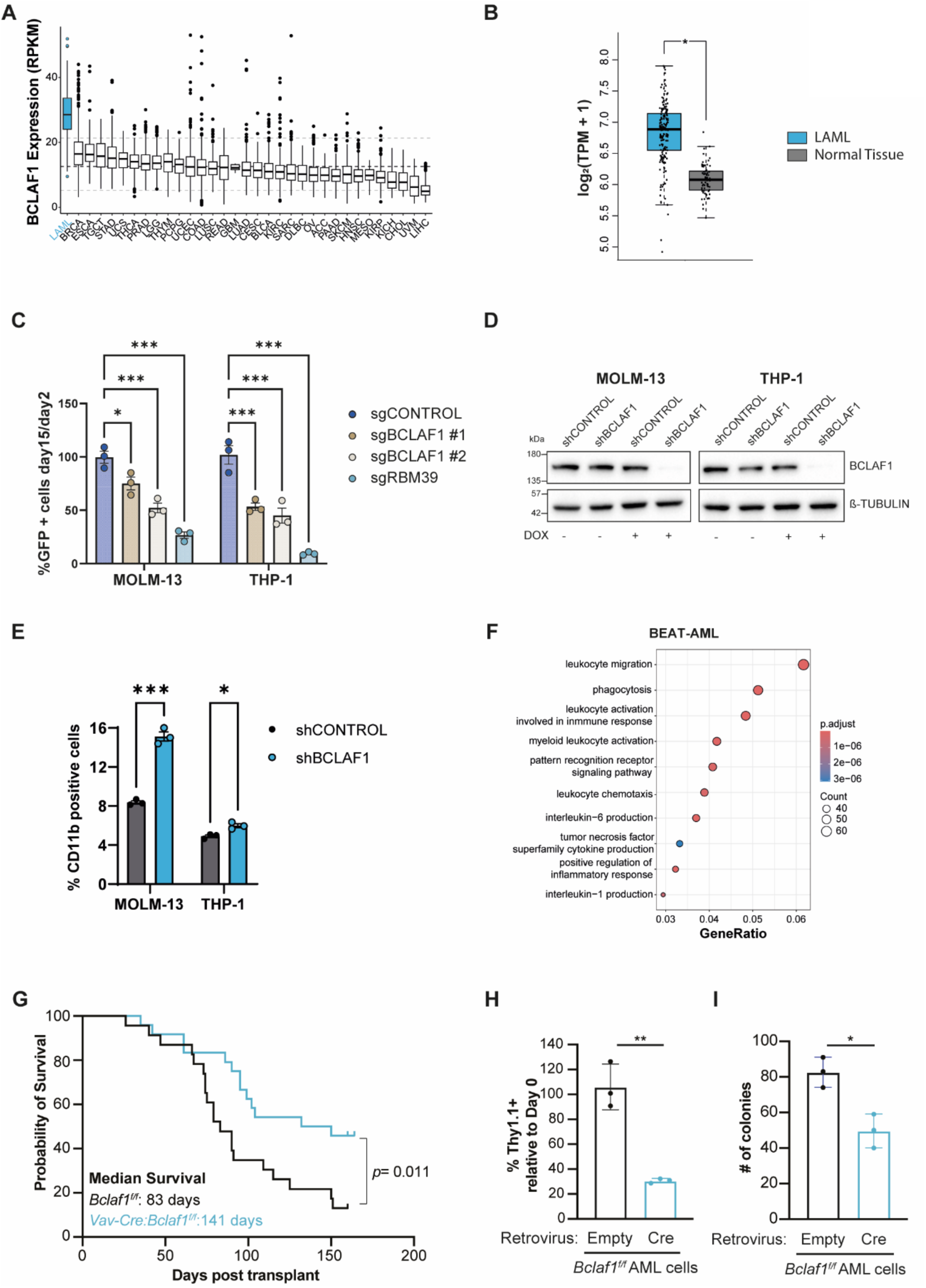
BCLAF1 supports AML pathogenesis. **A.** BCLAF1 expression levels across cancer types from the TCGA dataset. **B.** BCLAF1 expression in LAML patient samples (n = 173) and normal hematopoietic tissues (n = 70), shown as log_2_(TPM + 1). **C.** Growth competition experiments showing the percentage of GFP-positive cells between day 2 and day 15 following infection with LentiCRISPRv2GFP constructs expressing individual sgRNAs targeting the indicated genes. Data represent mean ± SEM of three biological replicates (*p < 0.05, ***p < 0.001; two-way ANOVA). **D.** Immunoblot analysis of BCLAF1 protein levels in shCONTROL and shBCLAF1 cells treated or not with DOX for 96 hours. β-tubulin was used as a loading control. **E.** Flow cytometry analysis showing the percentage of CD11b-positive cells in shCONTROL and shBCLAF1 cells treated with DOX for 96h. Data represent mean ± SEM of three biological replicates (*p < 0.05, ***p < 0.001; unpaired t-test). **F.** Dot plot of GO terms enriched among genes up-regulated in BCLAF1-low versus BCLAF1-high patient samples from the BEAT-AML cohort. GeneRatio indicates the proportion of significantly upregulated genes associated with each GO term. Dot size reflects the number of genes per term; color denotes adjusted p-value. **G.** Kaplan-Meier curve shows survival of mice transplanted with MLL-AF9 expressing *Bclaf1^f/f^*(black) or *Vav-Cre:Bclaf1^f/f^* (blue) HSPCs. Briefly, HSPCs from CD45.2^+^ *Bclaf1^f/f^* or *Vav-Cre:Bclaf1^f/f^* mice were transduced with retrovirus co-expressing MLL-AF9 and GFP then transplanted with 300,000 CD45.1^+^ recipient bone marrow cells into lethally-irradiated CD45.1^+^ recipient mice (schematic in Figure S1G). Leukemic burden was assessed by percentage of GFP^+^ cells in peripheral blood. Mice were euthanized once peripheral blood had ≥ 80% leukemia. Inset shows median survival. *Bclaf1^f/f^*, n=23; *Vav-Cre:Bclaf1^f/f^*, n=24. **H.** Primary *Bclaf1^f/f^* AMLs isolated from mice in G were transduced with retroviral vector expressing Thy1.1 (empty control, black) or Thy1.1 and Cre recombinase (blue) then expanded as mixed co-cultures with nontransduced cells. Bar graphs show percentage of transduced cells (Thy1.1^+^) at day 7 of mixed co-culture relative to day 0 transduction efficiency. Data repesent mean ± SD of 3 independent experiments (**p ≤ 0.01; unpaired t-test). **I.** Primary *Bclaf1^f/f^* AMLs isolated from mice in G were transduced with retroviral vectors as in D. Thy1.1^+^ cells were enriched by flow cytometric sorting and cultured in complete methocult media. Total colony forming units were quantitated at day 7 of culture. Data repesent mean ± SD of 3 independent experiments (*p ≤ 0.05; unpaired t-test).

To investigate the direct effects of BCLAF1 depletion, we generated MOLM-13 and THP-1 stable cell lines expressing a doxycycline (DOX)-inducible shRNA targeting *BCLAF1* (shBCLAF1) or a non-targeting control (shCONTROL). Upon DOX treatment, both cell lines showed efficient BCLAF1 depletion (Figure 1D) and reduced proliferation (Figure S1A), confirming a functional requirement for BCLAF1 in AML cells. To explore the underlying cause of this growth defect, we assessed the surface expression of CD11b, a marker of myeloid differentiation, and apoptosis levels using Annexin V staining. We found that depletion of BCLAF1 promotes AML cell differentiation, as indicated by increased CD11b expression in both MOLM-13 and THP-1 cells (Figure 1E). Comparatively, only modest changes in apoptosis were observed (Figure S1B). Similar results were obtained using a second independent shRNA targeting *BCLAF1* (Figure S1C-E), suggesting that BCLAF1 is required to maintain the undifferentiated leukemic state.

To assess whether BCLAF1 expression correlates with leukemic states in AML patients, we stratified BEAT-AML (28) cohort samples based on *BCLAF1* mRNA expression levels and defined the top 25% (145/578) as *BCLAF*1-high cases, and the bottom 25% (145/578) as *BCLAF1*-low cases. In total, we identified 3968 differentially expressed genes (false discovery rate (FDR) ≤ 0.05, foldchange (FC) ≥ 2) between these sets of samples (Table S1), including 1527 up-regulated transcripts in *BCLAF1*-low cases. Gene ontology analysis of up-regulated genes in *BCLAF1*-low patients showed an enrichment of categories such as leukocyte migration (GO:0050900) and myeloid leukocyte activation (GO:0002274) (Figure 1F), which are consistent with a more differentiated myeloid state. Notably, similar results were observed in the pediatric TARGET-AML cohort (Figure S1F and Table S2), further supporting the notion that BCLAF1 expression is associated with the maintenance of an undifferentiated leukemic transcriptional program in primary AML.

Next, we sought to determine whether loss BCLAF1 impairs of AML pathogenesis *in vivo*. Retroviral expression of the MLL-AF9 fusion gene in murine hematopoietic stem and progenitor cells (HSPCs) results in an established, aggressive form of AML.(29,30) We used this model of primary AML to assess the role of BCLAF1 in leukemic initiation *in vivo*. HSPCs from adult (8-week-old) donor CD45.2^+^ wild-type *Bclaf1^f/f^*or *Vav-Cre:Bclaf1^f/f^* mice, which have selective deletion of *Bclaf1* in hematopoietic cells,(26) were transduced with retrovirus co-expressing an MLL-AF9 fusion construct and a GFP reporter, then transplanted into lethally-irradiated CD45.1^+^ recipient mice (Figure S1G). Leukemic burden was monitored by percentage of GFP^+^ cells in peripheral blood. Of the mice that received MLL-AF9-transduced wild-type *Bclaf1^f/f^* HSPCs, 87% (20 of 23) developed AML with a median survival of 83 days (Figure 1G and S1H). In contrast, only 54% (13 of 24) of mice receiving MLL-AF9-transduced *Vav-Cre:Bclaf1^f/f^*HSPCs developed leukemia with a median survival of 141 days (Figure 1G). The surviving mice that received MLL-AF9-transduced *Vav-Cre:Bclaf1^f/f^*HSPCs had no evidence of AML (defined by absence of GFP+ cells) in bone marrow and spleen at 5 months post-transplant (Figure S1I). These results indicate that loss of BCLAF1 impairs AML initiation *in vivo*, as evidenced by a reduced incidence of leukemic transformation and prolonged survival in mice receiving *Bclaf1*-deficient MLL-AF9-expressing HSPCs.

To further investigate whether loss of BCLAF1 compromises AML expansion capacity, we cultured wild-type MLL-AF9-expressing *Bclaf1^f/f^* leukemias *in vitro* and transduced them with retrovirus expressing Cre recombinase to inactivate the *Bclaf1* allele. Deletion of *Bclaf1* led to impaired expansion of AML cells, as indicated by a more significant loss of cells expressing Cre recombinase (Thy1.1^+^ cells) compared with control cells expressing empty viral vector (Figure 1H). Additionally, sorted Cre-expressing *Bclaf1^f/f^* AML cells generated fewer colonies compared to sorted control *Bclaf1*-sufficient AML cells (Figure 1I). Altogether, our findings support an important role for BCLAF1 in AML pathogenesis.

### BCLAF1 regulates mRNA splicing in AML cells

BCLAF1 is a multifunctional protein implicated in diverse biological processes. (31–36) To gain mechanistic insight into BCLAF1 function in AML, we sought to identify its interacting partners by performing immunoprecipitation followed by mass spectrometry (IP-MS) in MOLM-13 cells. To enable specific pulldown of the protein, we transduced cells with a lentiviral construct expressing an ALFA-tagged (37) version of BCLAF1. Immunoblot analysis confirmed that ALFA-BCLAF1 was expressed at levels comparable to endogenous protein, ruling out potential overexpression artifacts (Figure S2A). IP-MS analysis identified 19 BCLAF1-specific interactors (Figure 2A and Table S3), which form a highly connected protein network enriched in components of the spliceosome complex (Figure S2B). Consistently, gene ontology analysis of BCLAF1-interactors indicated a strong enrichment for mRNA splicing and processing categories (Figure 2B), supporting a functional interaction between BCLAF1 and the splicing machinery in AML cells.

**Figure 2.**
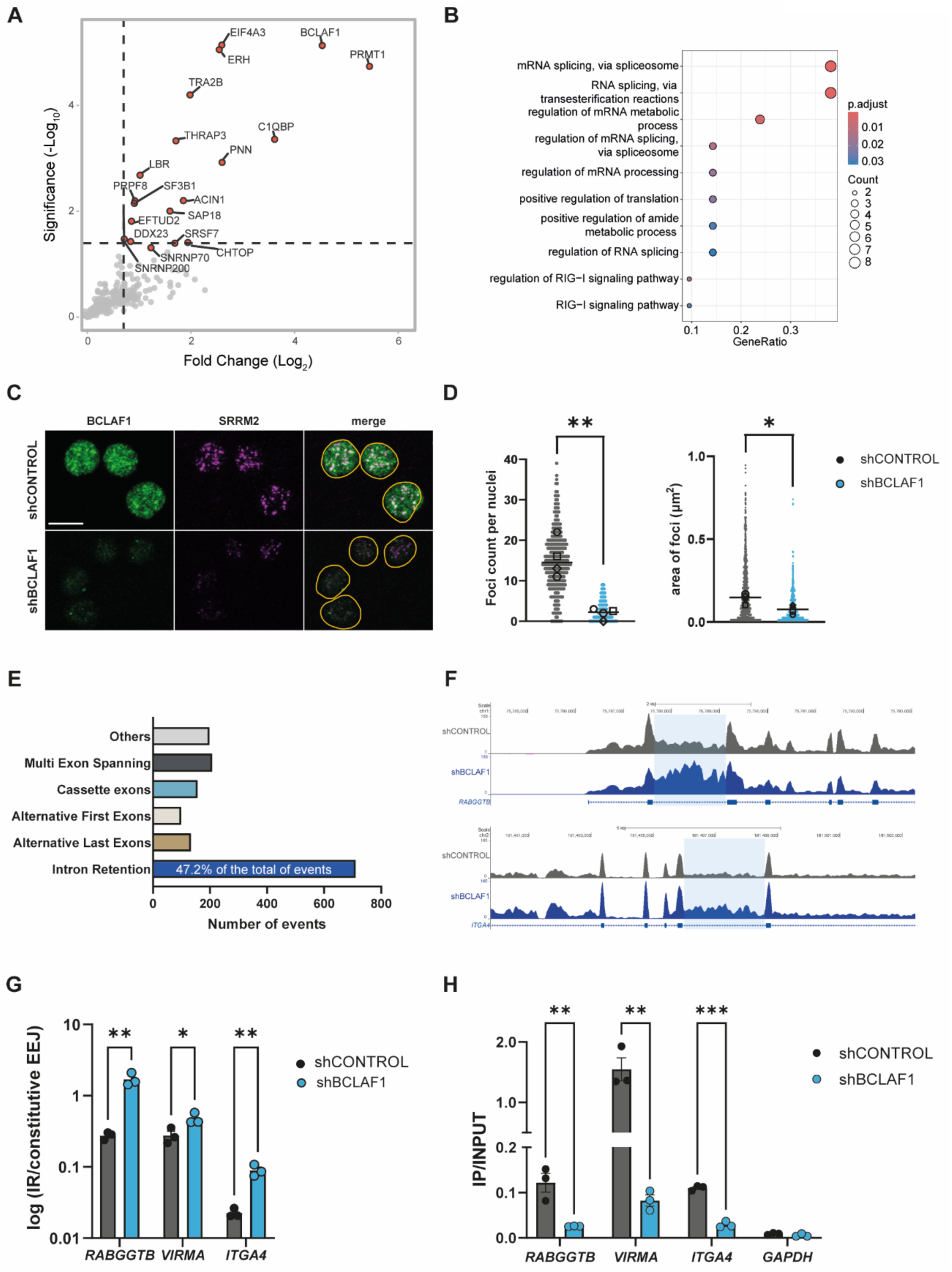
BCLAF1 regulates mRNA splicing in AML cells. **A.** Volcano plot showing BCLAF1-interacting proteins identified by immunoprecipitation followed by mass spectrometry (IP-MS). The x-axis indicates log2 fold change (ALFA-BCLAF1 IP vs. control), and the y-axis shows –log10 p-value. Significant interactors (fold change ≥ 1.5, p-value ≤ 0.05) are highlighted in red. **B.** Dot plot of GO terms enriched among BCLAF1 interactors. **C.** Immunofluorescence analysis of BCLAF1 and the nuclear speckle marker SRRM2 in shCONTROL and shBCLAF1 cells treated with DOX for 96h. Yellow lines mark the nuclear boundary (FIJI mask). Scale bar, 10 μm. **D.** Quantification of SSRM2 foci size and number per nucleus in shCONTROL and shBCLAF1 cells treated with DOX for 96 hours. Data represent the median values from four independent biological replicates, each shown as a distinct symbol. Statistical significance was assessed using the mean of medians and a Mann-Whitney test (*p < 0.05; **p < 0.01). **E.** Bar graph showing the distribution of alternative splicing event types significantly altered upon BCLAF1 depletion (96 hours of DOX treatment) as quantified by MAJIQ. **F.** Representative genome browser snapshots of RNA-seq data from shCONTROL and shBCLAF1 cells treated with DOX for 96 hours. **G.** RT-qPCR experiments showing relative intron retention (IR) levels of the indicated transcripts in shCONTROL and shBCLAF1 cells treated with DOX for 96 hours. Data represent mean ± SEM of three biological replicates (*p < 0.05, **p < 0.01; unpaired t-test). **H.** RIP-qPCR experiments showing BCLAF1 association with the indicated transcripts in shCONTROL and shBCLAF1 cells treated with DOX for 96 hours. *GAPDH* was used as a negative control. Data represent mean ± SEM of three biological replicates (**p < 0.01, ***p < 0.001; unpaired t-test).

Given that BCLAF1 cellular distribution varies across cancer types (31–34), we next examined its localization in MOLM-13 cells by immunofluorescence analysis. We observed that endogenous BCLAF1 localizes to dot-like structures throughout the nucleus that are reminiscent of nuclear speckles (Figure 2C), subnuclear compartments enriched in splicing factors.(38) Co-immunostaining with the nuclear speckle marker SRRM2 (39) confirmed partial colocalization (Figure 2C), indicating that BCLAF1 is associated with nuclear speckles in AML cells. Interestingly, depletion of BCLAF1 reduced both the size and number of nuclear speckles (Figure 2C and 2D), suggesting that BCLAF1 contributes to nuclear speckle integrity. In support of this, gene set enrichment analysis identified a marked downregulation of RNA splicing related genes in *BCLAF1*-low patients from both the adult BEAT-AML (Figure S2C) and the pediatric TARGET-AML (Figure S2D) cohorts, indicating a link between BCLAF1 expression and splicing-related transcriptional programs in primary AML. Taken together, these findings suggest that BCLAF1 associates with the spliceosome machinery and contributes to its proper function in AML cells.

The association of BCLAF1 with the spliceosome in AML cells prompted us to investigate its potential role in regulating mRNA splicing. We evaluated changes in alternative splicing by RNA sequencing (RNA-seq) of MOLM-13 cells upon depletion of BCLAF1. Using the quantification tool MAJIQ,(40) we found 1503 significantly altered splicing events (FDR ≤ 0.1, Δ percent spliced in (PSI) ≥ 0.05) (Table S4). The most frequently affected category was intron retention, accounting for nearly half of the total events (Figure 2E and 2F). Isoform-specific RT-qPCR validated BCLAF1-dependent splicing alterations of selected targets (Figure 2G), and similar results were obtained using a second shRNA targeting *BCLAF1* (Figure S2E). Altogether, these findings indicate that loss of BCLAF1 leads to widespread alterations in mRNA splicing.

To assess whether BCLAF1 directly associates with target mRNAs that undergo splicing changes, we performed BCLAF1 RNA immunoprecipitation sequencing (RIP-seq) experiments in MOLM-13 cells and identified 3567 significantly enriched transcripts in the BCLAF1-IP compared to input control (FDR ≤ 0.05, FC ≥ 1.5) (Table S5). Gene ontology analysis of BCLAF1-bound transcripts showed an enrichment for pathways related to mitotic and DNA double-strand break repair processes (Figure S2F), consistent with previously described roles of BCLAF1 in maintaining genome stability and cell cycle progression.(35,36,41) We also found an enrichment for mRNA splicing related categories (Figure S2F), reinforcing the notion that BCLAF1 contributes to the maintenance of the splicing machinery in AML cells. Interestingly, integration of the RIP-seq and RNA-seq datasets indicated a statistically significant overlap between transcripts bound by BCLAF1 and those exhibiting splicing alterations upon BCLAF1 knockdown (Figure S2G). This suggests that BCLAF1 controls alternative splicing, at least in part, through specific association with pre-mRNA targets. We validated a subset of these interactions by performing RIP-qPCR experiments, confirming both the specific association of BCLAF1 to selected transcripts and the loss of enrichment upon its depletion (Figure 2H). Collectively, these results indicate that BCLAF1 regulates mRNA splicing in AML cells, which is consistent with its association with the spliceosome complex and localization to nuclear speckles.

### BCLAF1 promotes amino acid metabolic gene expression

Our findings are consistent with a model in which BCLAF1 supports leukemic cell fitness through modulation of mRNA splicing. To further dissect the molecular pathways underlying its role in AML, we performed differential gene expression analysis in MOLM-13 cells following depletion of BCLAF1. This analysis identified 539 differentially expressed genes (FDR ≤ 0.05, FC ≥ 2) between shBCLAF1 and shCONTROL cells, including 226 up-regulated and 313 down-regulated transcripts (Figure 3A and Table S6).

**Figure 3.**
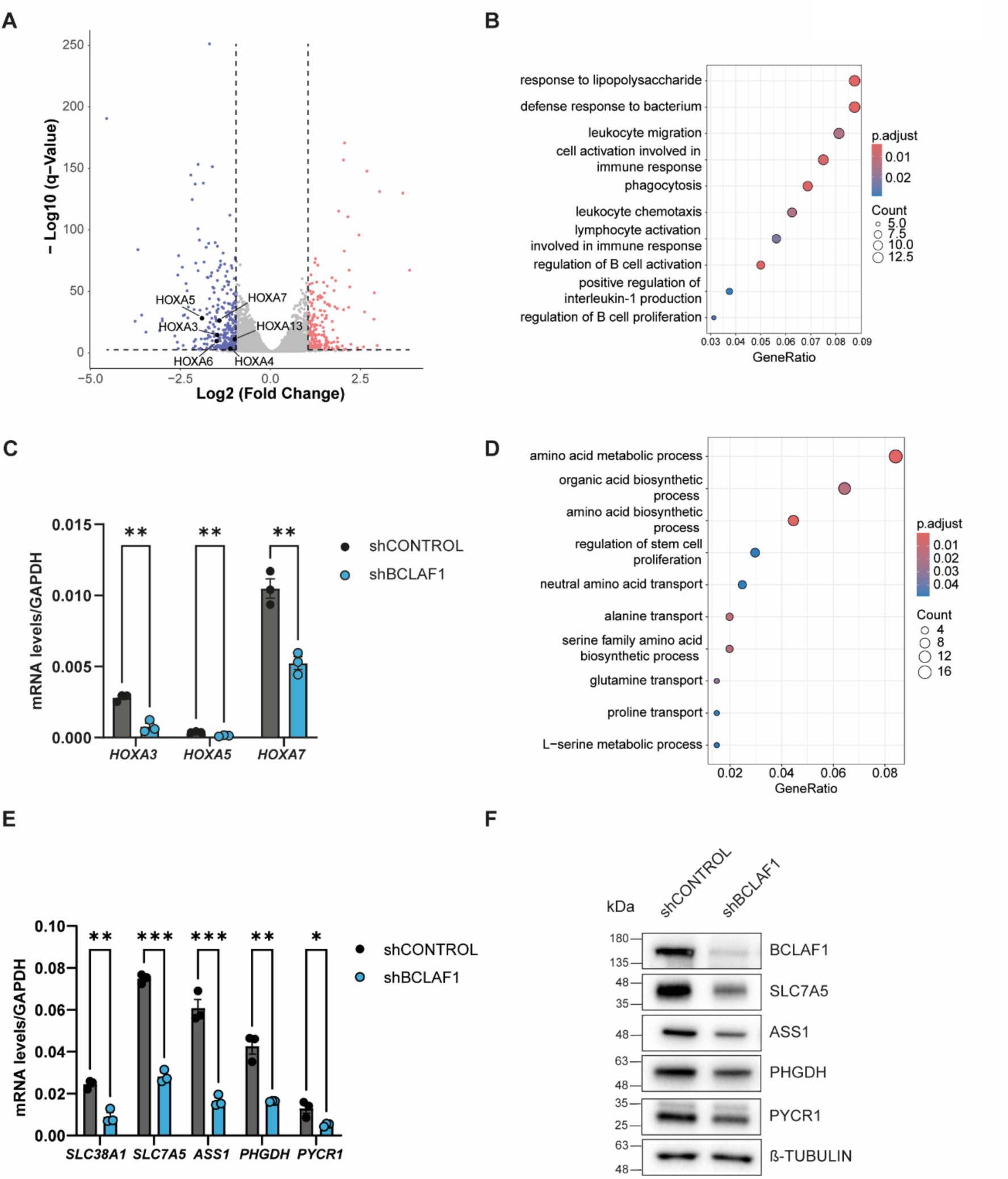
BCLAF1 promotes amino acid metabolic gene expression. **A.** Volcano plot showing differentially expressed genes in shBCLAF1 versus shCONTROL cells treated with DOX for 96 hours. The x-axis indicates log2 fold change and the y-axis represents –log10 adjusted p-value. Significantly up-regulated (red) and down-regulated (blue) genes are shown (fold change ≥ 2, adjusted p ≤ 0.05). **B.** Dot plot of GO terms enriched among genes up-regulated upon BCLAF1 depletion. **C.** RT-qPCR experiments showing relative mRNA levels of the indicated transcripts in shCONTROL and shBCLAF1 cells treated with DOX for 96 hours. Data were normalized to *GAPDH* and represent mean ± SEM of three biological replicates (**p < 0.01; unpaired t-test). **D.** Dot plot of GO terms enriched among genes down-regulated upon BCLAF1 depletion. **E.** RT-qPCR experiments showing relative mRNA levels of the indicated transcripts in shCONTROL and shBCLAF1 cells treated with DOX for 96 hours. Data were normalized to *GAPDH* and represent mean ± SEM of three biological replicates (*p < 0.05, **p < 0.01, ***p < 0.001; unpaired t-test). **F.** Immunoblot analysis of BCLAF1, SLC7A5, ASS1, PHGDH and PYCR1 protein levels in shCONTROL and shBCLAF1 cells treated or not with DOX for 96 hours. β-tubulin was used as a loading control.

Gene ontology analysis of up-regulated transcripts showed strong enrichment for immune-related processes such as leukocyte migration (GO:0050900) and cell activation involved in immune response (GO:0002263) (Figure 3B), suggesting a shift toward a more differentiated myeloid-like state. This is consistent with the observed increase in CD11b-positive cells upon BCLAF1 depletion (Figure 1D) and further supports a role for BCLAF1 in maintaining the undifferentiated state of AML cells. In line with this, we observed a marked downregulation of multiple *HOXA* cluster transcripts (Figure 3A), which are key regulators of leukemic transformation.(42) RT-qPCR analysis confirmed reduced expression of selected *HOXA* mRNAs upon BCLAF1 depletion (Figure 3C), and similar results were obtained using a second shRNA targeting BCLAF1 (Figure S3A).

We next focused on the subset of transcripts that were down-regulated upon BCLAF1 depletion. Gene ontology analysis showed a marked enrichment for pathways related to amino acid metabolism, particularly transport and biosynthesis (Figure 3D). These results were confirmed by RT-qPCR analysis, showing downregulation of representative genes associated with amino acid transport (*SLC38A1*, *SLC7A5*) and biosynthesis (*ASS1*, *PHGDH, PYCR1*) (Figure 3E), and similar results were obtained using a second independent shRNA targeting BCLAF1 (Figure S3B). In line with these transcriptional changes, immunoblot analysis confirmed reduced steady-state protein levels of SLC7A5, ASS1, PHGDH and PYCR upon BCLAF1 depletion (Figure 3F), further supporting the notion that BCLAF1 loss compromises amino acid metabolic capacity at both the transcript and protein levels.

Collectively, our findings establish the downregulation of amino acid metabolic genes as a major downstream consequence of BCLAF1 loss. Given the crucial role of amino acid metabolism in supporting leukemic cell growth and survival,(15–18,43) this may represent one mechanism through which BCLAF1 supports the undifferentiated leukemic state.

### Loss of BCLAF1 disrupts *de novo* amino acid synthesis

Based on our differential gene expression analysis, we next asked whether BCLAF1-dependent transcriptional changes translated into functional alterations in amino acid metabolism. We first assessed steady-state intracellular amino acid levels, which reflect the combined effects of synthesis, import, and catabolism. Gas chromatography–mass spectrometry (GC–MS) analysis revealed that while the total abundance of most detected amino acids was largely preserved upon BCLAF1 depletion, intracellular levels of proline and arginine were significantly reduced (Figure S4A and Table S7), indicating selective perturbations in amino acid homeostasis.

To directly assess *de novo* amino acid synthesis, we performed isotope tracing experiments in glucose-free media supplemented with a glucose isotopologue containing six heavy carbons (U-13C-glucose) for 24 hours. Labeling of upstream glycolytic intermediates, including glyceraldehyde-3-phosphate and pyruvate, as well as TCA cycle intermediates such as a-ketoglutarate, citrate and succinate, was comparable between shCONTROL and shBCLAF1 cells (Figure 4A and Figure S4B-C), indicating that central carbon metabolism remained intact. Consistent with this, steady-state levels of TCA-derived amino acids, including aspartate and glutamate, were not significantly altered upon BCLAF1 depletion (Figure S4D). In contrast, BCLAF1 knockdown cells displayed a marked reduction in the incorporation of ¹³C from glucose into serine, as evidenced by a significantly lower proportion of labeled serine isotopologues (the sum of M+1, M+2 and M+3, red) (4.29% vs. 15.45%, p < 0.05), accompanied by an accumulation of unlabeled serine (M+0, grey) (Figure 4B). This defect propagated downstream, resulting in significantly diminished labeling of glycine (2.57% vs. 7.2%, p < 0.01) and methionine (3.26% vs. 5.77%, p < 0.05) (Figure 4C-D). Despite preserved glycolytic flux, alanine labeling was also significantly reduced (68.01% vs. 87.65%, p < 0.001) (Figure 4E), indicating that defects in *de novo* amino acid synthesis extend beyond the serine–glycine pathway. Consistent with these metabolic alterations, expression of key biosynthetic enzymes involved in serine production (PHGDH, PSAT1, and PSPH), serine-to-glycine conversion (SHMT2), and alanine synthesis (GPT2) was markedly reduced upon BCLAF1 knockdown (Figure 4F), providing a direct mechanistic basis for the observed defects. Collectively, these findings suggest that BCLAF1 loss selectively disrupts *de novo* amino acid synthesis through diminished expression of its associated metabolic enzymes.

**Figure 4.**
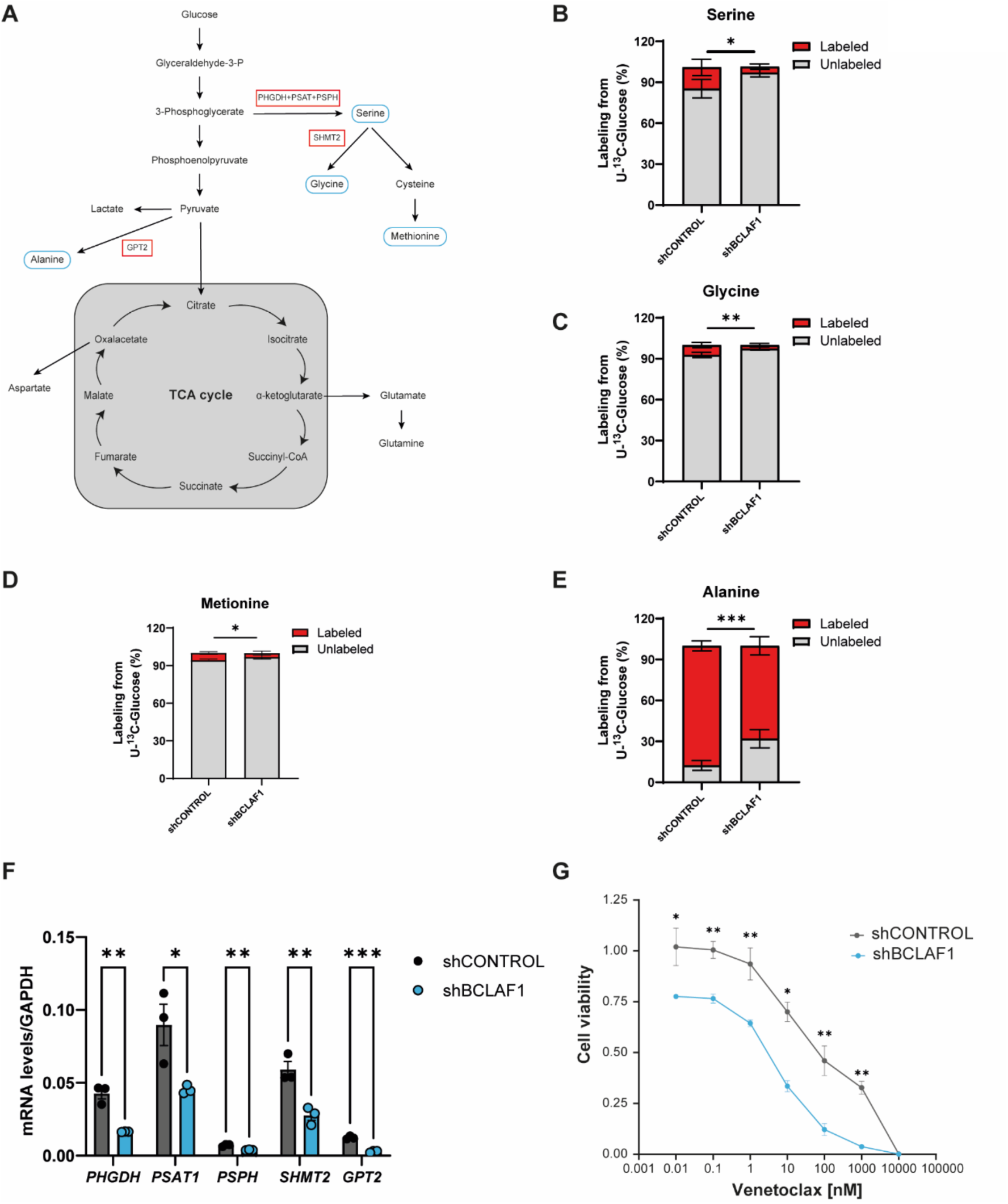
Loss of BCLAF1 disrupts *de novo* amino acid synthesis. **A.** Schematic of amino acid biosynthesis pathways. **B. C. D.** and **E.** Percentage of unlabeled (M+0; gray) or labeled (the sum of M+1, M+2, and M+3; red) amino acid levels. Data are shown as mean ± SEM of four technical replicates from one representative experiment (*p < 0.05, **p < 0.01, ***p < 0.001; two-way ANOVA). The experiment was independently repeated with similar results. **F.** RT-qPCR experiments showing relative mRNA levels of the indicated transcripts in shCONTROL and shBCLAF1 cells treated with DOX for 96 hours. Data were normalized to *GAPDH* and represent mean ± SEM of three biological replicates (*p < 0.05, **p < 0.01, ***p < 0.001; unpaired t-test). **G.** Cell viability assay (CellTiter-Blue-based) in shCONTROL and shBCLAF1 cells treated or not with DOX for 96 hours, followed by 96 additional hours of Venetoclax treatment. Data is normalized to control (DMSO) and represent mean ± SEM of four biological replicates (*p < 0.05, **p < 0.01; unpaired t-test).

Amino acid metabolism plays a central role in AML maintenance. (44) Thus, we next asked whether BCLAF1 depletion could impact cell sensitivity to venetoclax, a BCL-2 inhibitor shown to impair amino acid uptake and mitochondrial metabolism in LSCs. (15) We treated shCONTROL and shBCLAF1 MOLM-13 cells with increasing concentrations of venetoclax 96 hours after doxycycline induction. BCLAF1-depleted cells showed significantly increased sensitivity to venetoclax, consistent with a compromised metabolic state (Figure 4G). These results suggest that BCLAF1 loss enhances sensitivity to venetoclax, highlighting a potential therapeutic vulnerability in AML.

### BCLAF1 supports amino acid metabolism through ATF4

Our transcriptomic analyses in MOLM-13 cells indicated that depletion of BCLAF1 leads to widespread changes in both gene expression and mRNA splicing. However, integration of differential expression and alternative splicing analyses revealed limited overlap (Figure S5A), suggesting that most BCLAF1-dependent splicing alterations do not directly impact global steady-state mRNA abundance in AML cells.

To better understand how BCLAF1 depletion drives transcriptional changes, we performed an unbiased transcription factor enrichment analysis using the TRRUST database.(45) This analysis identified ATF4, a key transcriptional regulator of amino acid metabolism,(46) as the most significantly enriched factor among the BCLAF1-downregulated gene set (Table S8). Supporting this connection, analysis of ENCODE ChIP-seq data for ATF4 in K562 AML cells revealed that 75% (18 of 24) of BCLAF1-downregulated genes annotated in amino-acid metabolic pathways display ATF4 binding (Figure S5B and S5C). These data suggest that the transcriptional effects of BCLAF1 depletion may be mediated, at least in part, through reduced ATF4 activity. In line with this hypothesis, immunoblot analysis of MOLM-13 cells showed a marked reduction in ATF4 protein levels upon BCLAF1 depletion (Figure 5A). Likewise, MLL-AF9 leukemias derived from *Bclaf1*-deficient *Vav-Cre:Bclaf1^f/f^* mice exhibited decreased ATF4 protein levels compared to leukemias derived from wild-type *Bclaf1^f/f^*controls (Figure 5B), suggesting that BCLAF1 supports ATF4 expression in primary AML cells.

**Figure 5.**
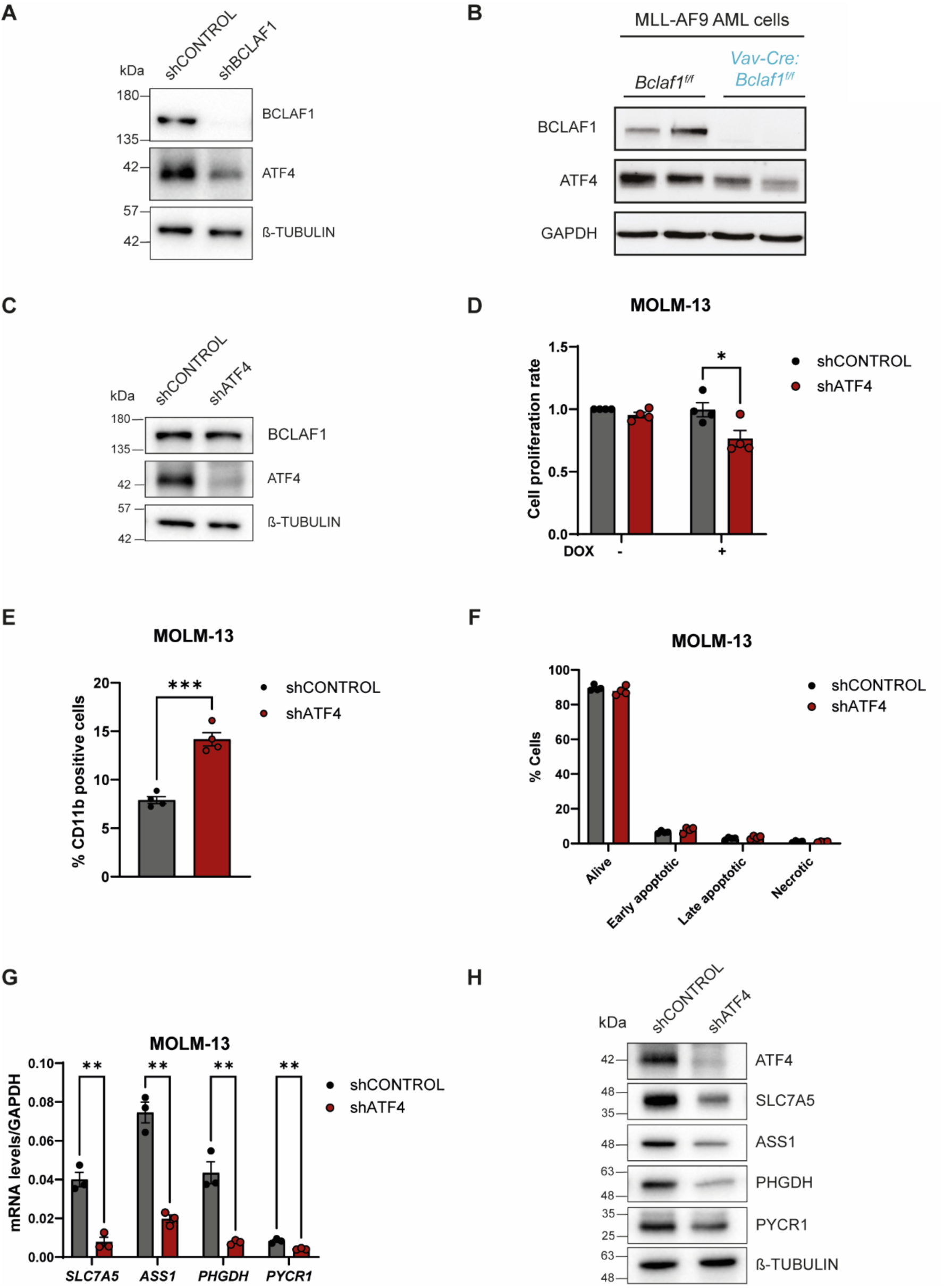
BCLAF1 supports amino acid metabolism through ATF4. **A.** Immunoblot analysis of BCLAF1 and ATF4 protein levels in shCONTROL and shBCLAF1 cells treated with DOX for 96 hours. β-tubulin was used as a loading control. **B.** Immunoblot analysis of BCLAF1 and ATF4 protein levels in MLL-AF9-expressing AML cells of indicated genotypes. GAPDH was used as a loading control. **C.** Immunoblot analysis of BCLAF1 and ATF4 protein levels in shCONTROL and shATF4 cells treated with DOX for 96 hours. β-tubulin was used as a loading control. **D.** Cell proliferation assay (CellTiter-Blue-based) in shCONTROL and shATF4 cells treated or not with DOX for 96 hours, followed by 96 additional hours in culture. Data represent mean ± SEM of three biological replicates (*p < 0.01; paired t-test). **E.** Flow cytometry analysis showing the percentage of CD11b-positive cells in shCONTROL and shATF4 cells treated with DOX for 96h. Data represent mean ± SEM of four biological replicates (***p < 0.001; unpaired t-test). **F.** Annexin V/PI flow cytometry analysis showing the distribution of live, early apoptotic, late apoptotic, and necrotic cells following 96h of DOX treatment. Data represent mean ± SEM of three biological replicates (two-way ANOVA). **G.** RT-qPCR experiments showing relative mRNA levels of the indicated transcripts in shCONTROL and shATF4 cells treated with DOX for 96 hours. Data were normalized to *GAPDH* and represent mean ± SEM of three biological replicates (**p < 0.01; unpaired t-test). **H.** Immunoblot analysis of ATF4, SLC7A5, ASS1, PHGDH and PYCR1 protein levels in shCONTROL and shATF4 cells treated or not with DOX for 96 hours. β-tubulin was used as a loading control.

To determine whether this relationship between BCLAF1 and ATF4 expression levels extends beyond MLL-AF9-driven leukemias, we examined OCI-AML3 cells, which carry NPM1 and DNMT3A mutations. BCLAF1 depletion in OCI-AML3 cells also led to a pronounced reduction in ATF4 protein levels (Figure S5D), which was accompanied by downregulation of ATF4 target genes involved in amino acid metabolism (Figure S5E), impaired cell proliferation (Figure S5F) and increased sensitivity to venetoclax treatment (Figure S5G). These data indicate that the BCLAF1–ATF4 regulatory axis operates across genetically distinct AML subtypes.

ATF4 activity and target gene expression are known to be elevated in primary LSCs, (47) and several studies have implicated ATF4 in AML pathogenesis. (48–50) However, the precise molecular contribution of ATF4 to AML maintenance remains incompletely defined. To test whether ATF4 mediates the functional effects of BCLAF1 loss, we generated MOLM-13 cell lines expressing a DOX-inducible shRNA targeting *ATF4*. ATF4 depletion did not alter BCLAF1 protein levels (Figure 5C), consistent with BCLAF1 functioning upstream of ATF4. Notably, ATF4 knockdown phenocopied BCLAF1 depletion, resulting in impaired cell proliferation (Figure 5D) and increased CD11b expression (Figure 5E) with no evident effects in apoptosis (Figure 5F). Moreover, ATF4 knockdown also reduced the expression of amino acid metabolic genes downregulated in BCLAF1-deficient cells, both at the mRNA (Figure 5G) and protein (Figure 5H) levels. Taken together, these findings support a model in which BCLAF1 promotes amino acid metabolic programs important for AML maintenance through ATF4.

### BCLAF1 controls *ATF*4 mRNA alternative splicing

Next, we sought to determine how BCLAF1 regulates ATF4 expression. *ATF4* mRNA has two major splice variants: variant 1 (V1, ENST00000337304.2), which retains the first intron, and variant 2 (V2, ENST00000674920.3), which encodes the canonical ATF4 protein (Figure S6A). V1 is highly unstable in AML cells due to its nuclear retention and degradation, whereas V2 is efficiently exported and translated into functional ATF4 protein.(51) To investigate whether BCLAF1 regulates this splicing event, we performed isoform-specific RT-qPCR analysis. We first confirmed that V1 is expressed at substantially lower levels than V2 under basal conditions (Figure S6B), as previously reported.(51) Upon BCLAF1 depletion, we observed a significant increase in the V1/V2 ratio (Figure 6A), indicating a shift in splicing toward the intron-retaining isoform. This shift reflected both an increase in V1 levels and a concomitant reduction in V2 (Figure S6C), suggesting that BCLAF1 controls the balance between these isoforms. As a result, total *ATF4* mRNA levels were reduced in BCLAF1-depleted cells (Figure 6B), in line with the reported instability of the V1 isoform in AML cells.(51) These results indicate that BCLAF1 promotes the productive splicing of *ATF4* pre-mRNA to the stable, protein-coding V2 isoform, thereby sustaining *ATF4* transcript abundance.

**Figure 6.**
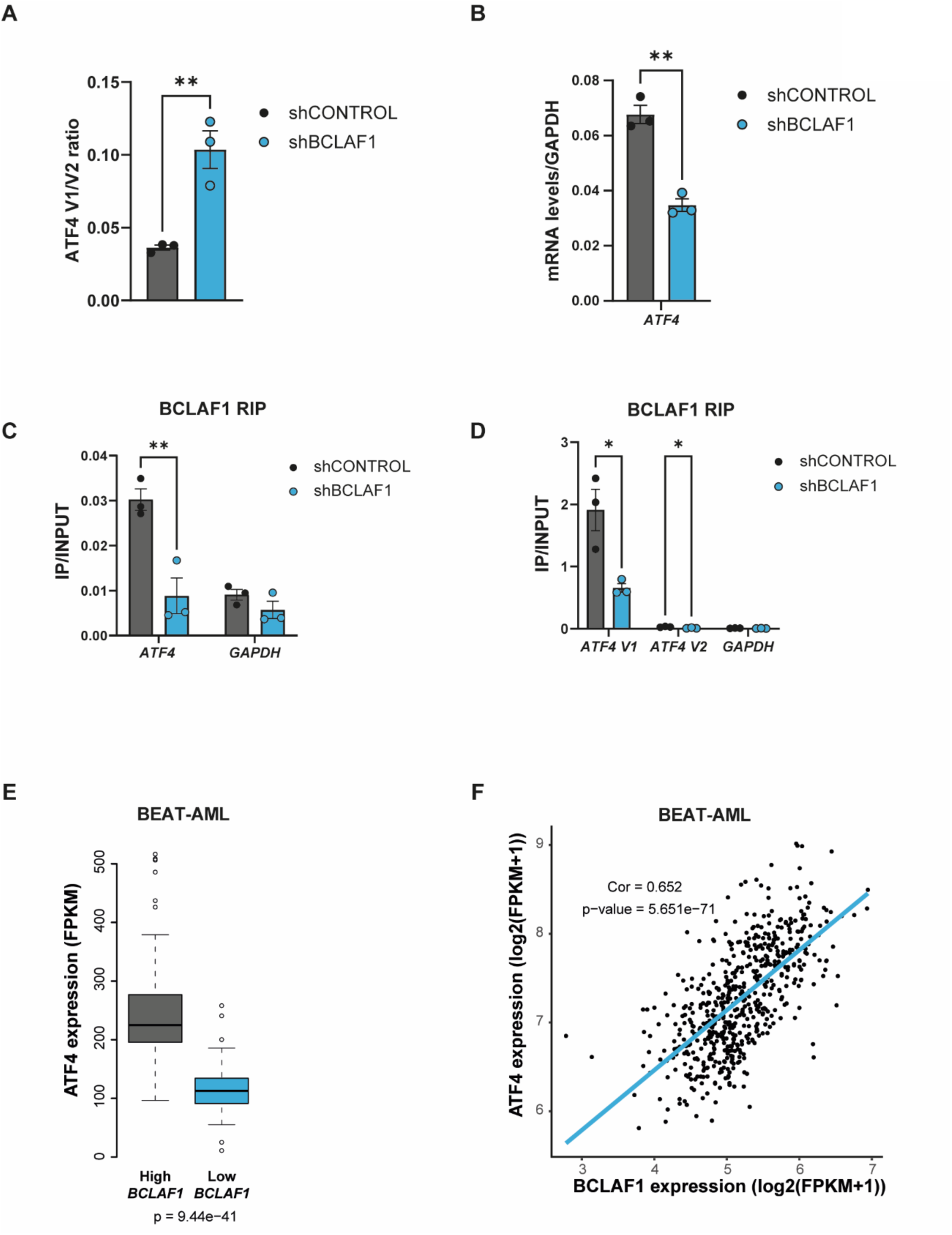
BCLAF1 controls *ATF4* mRNA alternative splicing. **A.** and **B.** RT-qPCR experiments showing relative mRNA levels of the indicated transcripts in shCONTROL and shBCLAF1 cells treated with DOX for 96 hours. Data were normalized to *GAPDH* and represent mean ± SEM of three biological replicates (**p < 0.01; unpaired t-test). **C.** and **D.** RIP-qPCR experiments showing BCLAF1 association with the indicated transcripts in shCONTROL and shBCLAF1 cells treated with DOX for 96 hours. *GAPDH* was used as a negative control. Data represent mean ± SEM of three biological replicates (*p < 0.05, **p < 0.01; unpaired t-test). **E.** Box plot showing *ATF4* mRNA levels in *BCLAF1*-high (top 25%) and *BCLAF1*-low (bottom 25%) patient samples from the BEAT-AML cohort. Statistical significance was assessed using Wilcoxon Rank Sum and Signed Rank Tests (one-sided). **F.** Scatter plot showing the correlation between *BCLAF1* and *ATF4* mRNA levels across BEAT-AML patient samples. Pearson correlation coefficient (r) and p-value are indicated.

Our RIP-seq analysis identified *ATF*4 mRNA as a prominent BCLAF1-bound transcript (Table S5). To validate this interaction, we performed RIP-qPCR experiments following BCLAF1 knockdown. We found that BCLAF1 specifically associates with the *ATF4* transcript, and that this interaction is reduced upon BCLAF1 depletion (Figure 6C). Moreover, this interaction was markedly enriched for the V1 isoform when compared with V2 (Figure 6D), suggesting preferential association of BCLAF1 with the unspliced pre-mRNA. This observation aligns with the presence of a conserved non-canonical weak splice donor site in intron 1,(51) and raises the possibility that BCLAF1 facilitates efficient splice site recognition, thereby promoting intron removal. Collectively, our data support a model in which BCLAF1 safeguards productive *ATF4* mRNA splicing by directly associating with its transcript. This regulatory step is required to sustain ATF4 protein levels and downstream transcriptional programs controlling amino acid metabolism in AML cells.

To validate the clinical relevance of these findings, we next analyzed RNA-seq data from the BEAT-AML cohort. Notably, we found that *BCLAF1*-low patients displayed significantly reduced *ATF4* mRNA levels compared to *BCLAF1*-high cases (Figure 6E). Consistent with this, plotting *BCLAF1* versus *ATF4* mRNA levels revealed a strong positive correlation across all patient samples (Figure 6F), indicating that this regulatory relationship extends to primary human AML. This association was also observed, although to a lesser extent, in the pediatric TARGET-AML cohort (Figure S6D–E), further reinforcing the potential clinical relevance of the BCLAF1–ATF4 axis. Taken together, these data indicate that BCLAF1 expression is closely linked to ATF4 levels in primary AML.

## Discussion

Multiple studies have identified distinct RNA-binding proteins (RBPs) as essential for leukemic cell survival, (27,52–56) underscoring the importance of post-transcriptional control in AML pathogenesis. Alternative mRNA splicing has been shown to alter the expression of key AML-associated genes independently of somatic mutations, thereby contributing to leukemogenesis and disease progression. (57,58) Moreover, dysregulation of splicing regulatory mechanisms has been implicated in therapy resistance. (59) Together, these findings position splicing regulation as a key contributor to AML evolution and therapeutic response. In this study, we identify BCLAF1 as an important post-transcriptional regulator in AML and show that it promotes leukemic maintenance by regulating alternative mRNA splicing, establishing it as a fundamental RBP dependency in AML.

BCLAF1 was originally identified as a BCL2-interacting protein with pro-apoptotic and transcriptional repressor functions.(60) Subsequent studies have implicated BCLAF1 in a broad range of biological processes, including DNA damage response, viral infection, and cellular differentiation.(61) More recently, BCLAF1 has emerged as an RNA-binding protein involved in mRNA metabolism.(35,36,41,62,63) Here, we show that BCLAF1 physically interacts with core spliceosome components and regulates hundreds of alternative splicing events in AML cells, with a predominant impact on intron retention. Our data indicate that these effects are mediated, at least in part, through its specific association with mRNA targets, supporting a model in which BCLAF1 functions as a splicing co-regulator. These findings provide mechanistic insight into previous observations linking BCLAF1 to AML pathogenesis (64,65) and further establish it as a central post-transcriptional regulator in leukemia.

We identify the stress-responsive transcription factor ATF4, a fundamental regulator of amino acid metabolism and cellular homeostasis,(46,66) as a key downstream effector of BCLAF1. BCLAF1 associates with *ATF4* mRNA and regulates its splicing to minimize an intron-retaining, unstable variant and maintain a stable, protein coding isoform. Both ATF4 activity and target gene expression have been shown to be elevated in primary LSCs (47), and multiple studies have implicated ATF4 in supporting AML pathogenesis. (48–50) Our findings position BCLAF1 as an upstream post-transcriptional regulator of ATF4, uncovering a mechanistic link by which splicing control contributes to amino acid metabolic adaptation and leukemic maintenance.

Recent studies have highlighted the therapeutic potential of targeting amino acid metabolic pathways in AML. (14,44) For example, reducing amino acid import through the combination of venetoclax and azacitidine selectively eliminates LSCs by impairing mitochondrial respiration.(15) Moreover, AML cells display distinct metabolic dependencies on specific amino acids, including on glutamine,(17,43) serine (18,21) and arginine.(19,20) However, leukemia cells can adapt to nutrient stress by inducing the expression of ATF4, (67) which could explain the limited efficacy of amino acid deprivation strategies when used as monotherapies.(68,69) Our results suggest that BCLAF1 may contribute to this adaptive capacity by regulating ATF4 expression, highlighting its potential as a combinatorial therapeutic target to enhance the efficacy of metabolic interventions in AML.

BCLAF1 is ubiquitously expressed but displays cell-type dependent functions.(61) During tumorigenesis, it can act as either a tumor promoter (32,33,70–73) or tumor suppressor (74–77) depending on the cancer type and cellular context. Interestingly, our previous work showed that although BCLAF1 promotes fetal HSC development, it is dispensable for the maintenance of adult HSCs under steady-state conditions.(26) This differential requirement raises the possibility that targeting BCLAF1 may selectively impair leukemia cells while sparing normal hematopoiesis, offering a potentially favorable therapeutic window.

## Methods

### Cell lines

Human tumor cell lines MOLM-13 (male; DSMZ, ACC-554; RRID CVCL_2119) and THP-1 (male; ATCC, TIB-202; RRID CVCL_0006) cells were cultured in RPMI 1640-Glutamax (61870036, GIBCO) supplemented with 10% FBS (16SV30160.03, CYTIVA) and 1% penicillin/streptomycin (5525-053-Cl, CYTIVA). OCI-AML3 (male; DSMZ, ACC 582; CVCL_1844) cells were cultured in MEM alpha modified with L-glutamine, Ribo-y Deoxyribonucleosides, (SH30265.01, CYTIVA) supplemented with 10% FBS (16SV30160.03, CYTIVA) and 1% penicillin/streptomycin (5525-053-Cl, CYTIVA). HEK293T cells (female; ATCC, CRL-3216; RRID: CVCL_0063) were cultured in DMEM High Glucose (D6546, Sigma-Aldrich) supplemented with 10% FBS (16SV30160.03, CYTIVA) and 1% penicillin/streptomycin/glutamine (5525-053-Cl, 5525-005-Cl, CYTIVA). All cells were grown at 37°C in a humidified incubator with 5% CO_2_. MOLM-13, THP-1 and OCI-AML3 cells were used as AML models; HEK293T for lentiviral production. Cell lines were regularly tested and confirmed negative for mycoplasma.

### Mice

*Bclaf1f/f* mice (RRID:MMRRC_046783-UCD) were previously described and were obtained from the Mutant Mouse Resource and Research Center (MMRRC) at University of California at Davis, an NIH-funded strain repository, and were donated to the MMRRC by The KOMP Repository, University of California, Davis. Vav-Cre mice (RRID:IMSR_JAX:008610) and Mx-Cre mice (RRID:IMSR_JAX:003556) were purchased from The Jackson Laboratory. CD45.1 mice (RRID:IMSR_CRL:494) mice were purchased from Charles River Laboratories. Mice were bred and housed according to IACUC guidelines at Washington University in St. Louis. Both sexes were used equivalently, except for female-only transplant recipients.

### Plasmids

pMSCV-IRES-GFP II (pMIG II) was a gift from Dario Vignali (Addgene plasmid #52107; RRID:Addgene_52107). pMIG-FLAG-*MLL-AF9* was a gift from Daisuke Nakada (Addgene plasmid #71443; RRID:Addgene_71443). Cre was cloned into pMSCV-IRES-Thy1.1 to create pMIT-Cre. pMSCV-IRES-Thy1.1 was a gift from Anjana Rao (Addgene plasmid #17442; RRID:Addgene_17442). Tet-pLKO-puro was a gift from Dmitri Wiederschain (Addgene plasmid # 21915; RRID:Addgene_21915), LentiCRISPRv2GFP was a gift from David Feldser (Addgene plasmid # 82416; RRID:Addgene_82416) and pHIV-Zsgreen was a gift from Bryan Welm & Zena Werb (Addgene plasmid # 18121; RRID:Addgene_18121). pVSV-G was a gift from Akitsu Hotta (Addgene plasmid # 138479; RRID:Addgene_138479). p8.91 was a gift from Simon Davis (Addgene plasmid # 187441; RRID:Addgene_187441).

### Virus production

For lentiviral production HEK293T cells were transfected with the appropriate lentiviral vector mix (PLKO.1 for shRNA, LentiCRISPR-v2GFP for CRISPR-Cas9 and pHIV-Zsgreen for overexpression) together with the packaging plasmids p8.91 and pVSVg at a 1:1.5:0.5 using Lipofectamine 2000 reagent (Thermo Fisher Scientific, 11668019). Specific oligos for each plasmid are indicated in Table S9. Supernatant was harvested 48 and 72 hours after transfection. For human cells transduction, cells and viral supernatants were mixed with 8 μg/ml (human) polybrene (TR-1003-G, Millipore), followed by spinfection (90 min, 850 *g*, RT) and further incubated overnight at 37 °C. Cells were collected on the following day. Retrovirus was produced using Phoenix Eco cells (ATCC Cat# CRL-3214, RRID:CVCL_H717). On day 1, cells were plated at 500,000 cells/well in a 6-well plate in DMEM media containing MEM non-essential amino acids, glutamine, sodium pyruvate, beta-mercaptoethanol, penicillin, streptomycin, and 10% fetal bovine serum. On day 2, cell media was changed to antibiotic free media. Plasmid DNA was mixed and incubated with lipofectamine (Invitrogen) before dropwise transfection according to manufacturer’s protocol. On day 3, lipofectamine-containing media was replaced with standard media. On day 4, virus-containing media was harvested from the supernatant of cells.

### MLL-AF9 leukemia studies

CD45.2^+^ *Bclaf1f/f* or *Vav-Cre:Bclaf1f/f* donor mice were injected intraperitoneally with 150 mg/kg 5-fluorouracil (5-FU) 6 days prior to cell collection. 5-FU (Sigma #F6627) was dissolved to 25 mg/ml in PBS and pH was adjusted to 9-10 with 10N NaOH. On day 6 (after 5-FU), bone marrow was collected from hind limbs, spine, and pelvis then enriched for c-Kit (CD117)-expressing cells using magnetic beads. Briefly, cells were incubated with biotinylated anti-c-Kit antibody (Biolegend #105803) for 20 minutes, then washed and incubated with MojoSort Streptavidin nanobeads (Biolegend #480016) for 15 minutes. Cells were washed then passed through magnetic MS columns (Miltenyi #130-042-201) according to manufacturer’s instructions. Enriched cells were resuspended at 2 x 10^6^ cells/mL in transduction media (Stemspan SFEM (StemCell Technologies), 100 ng/ml recombinant murine TPO, 100 ng/mL recombinant murine SCF (PeproTech), penicillin/streptomycin (Gibco)) and plated with polybrene (5 mg/mL, Sigma) and equivalent volumen (1:1) of viral supernatant in a 48-well plate. Cells were spinfected for 2 hours at room temperature at 1600 rpm. After spinfection, media was replaced with fresh transduction media with no polybrene, and cells were cultured overnight. The following morning, cells were spinfected again (as above) and cultured with fresh media for 6 hours before harvest and transplantation. For all transplants, CD45.1^+^ recipient mice received a split dose (≥ 3 hours apart) of lethal irradiation (11 Gy total). Primary recipients were administered 50,000–100,000 infected CD45.2^+^ donor cells with 300,000 CD45.1^+^ support cells. Cells were administered via retroorbital injection, and recipients were treated with trimethoprim/sulfamethoxazole for 14 days post-transplant. Mice were monitored weekly by peripheral blood GFP^+^ analysis and euthanized when peripheral blood showed ≥ 80% GFP positivity. Bone marrow and spleen were analyzed by flow cytometry for CD45.2^+^GFP^+^ blasts.

### *In vitro* murine AML studies

*Bclaf1f/f* MLL-AF9 cells harvested from transplanted mice as above were transduced with either pMSCV-IRES-Thy1.1 or pMCSV-IRES-Cre-Thy1.1 at 1500 rpm for 90 min, then cultured overnight in DMEM with 20% FBS and 10 ng/mL IL-3. Thy1.1^+^ cells were quantified by flow cytometry on day 7. For MIT-Cre CFU assays, Thy1.1^+^ sorted cells were plated in Methocult M3434 supplemented with IL-3. Colonies were counted at day 7.

### GFP-KO competition assay

GFP levels were measured by flow cytometry 2 days after infecting to set the starting baseline GFP percentage for each gRNA. The represented ratio was calculated using GFP^+^ cells day15 / GFP+ cells day2, which stands for the depletion of cells targeted for each guide. Two gRNAs were used to target BCLAF1, one for the Rosa26 locus as a negative control and one for RBM39 as a positive control. Flow cytometry data was analyzed using FlowJo software.

### Flow Cytometry

Cells were collected after 4 days of doxycycline induction of the MOLM-13 and THP-1 knock-down cell lines for BCLAF1 and control. These samples were stained using CD11b surface marker and Annexin V kit (Annexin V-FITC Kit, Miltenyi Biotec). In both cases, the procedure followed according to the manufacturer’s indications. Data was analyzed by using FACSCalibur and FlowJo. Mice samples were stained with antibodies for 20 min in staining media PBS with 2% FBS. Antibodies include CD11b-APC, CD19-PE-Cy7, GR1-PerCP-Cy5.5, CD3-PE, CD45.1-APC-Cy7, CD45.2-AF700 and Thy1.1-APC. Cells were washed and resuspended in staining media with DAPI to exclude dead cells. All flow cytometric analyses and sorting were performed on a BD Fortessa LSR or BD Aria Fusion.

### Western blots

Resolved proteins were transferred to a 0.45 µm PDVF membrane (IPVH00010, Millipore). Primary and secondary antibodies used are specified on Table S9.

### Proliferation assays

Knock-down cell lines (shCONTROL and shBCLAF1) were either untreated or treated with doxycycline (200 ng/mL) for 96h. Afterwards, cells were seeded in 96-well plates for another 96h using the corresponding treatments. Seeding concentration was 30.000 cells/mL and at least 3 wells per condition were included. Cell proliferation was measured using MTT assay (475989, Sigma). For drug sensitivity assays, cells were treated with a 6-point, ten-fold dilution series of Venetoclax (Bcl-2 inhibitor). Cells were treated for 96 hours with the drug, keeping the knock-down induction, and cell growth was measured using CellTiter-Blue® Fluorescence kit (G8081, Promega) following manufacturer’s indications. Fluorescence analysis for both assays was performed using the Varioskan Lux (ThermoFisher) plate reader.

### Immunofluorescence and confocal imaging

MOLM-13 cells were placed on a poly-L-lysinated (Sigma-Aldrich, P8920) coverslip and incubated for 30 min at RT in PBS. Cells were pre-extracted and then fixed using 4% formaldehyde for 10 min at RT, followed by a permeabilization with 0.2% PBS Triton X-100 for 2 min at RT. Cells were then blocked in 5% BSA in PBS for 30 min at RT. Primary antibodies were prepared in 1% BSA in PBS and incubated for overnight at 4°C in agitation. After three washes with PBS 0.1% Tween20, secondary antibody was prepared in PBS 1% BSA and added for 30 min at RT protected from light. DAPI was then incubated for 1 min at RT after 3 new PBS 0.1% Tween20 washes. Finally, coverslips were washed with PBS, dried and placed onto microscope slides using VectaShield mounting medium. Primary and secondary antibodies are specified on the Supplementary Material Table. Images were captured using SP5 confocal (Leica) with a HCX PL APO lambda blue 63x/1.40 OIL UV objective. The number of SC35-positive nuclear speckles per nucleus and the area of SC35-positive nuclear speckles were automatically quantified from 4 biological replicates using a custom ImageJ plug-in. List of antibodies in Table S9.

### RT-qPCR

Total RNA was purified using either the Direct-zol RNA MiniPrep Kit (Zymo Research, R2050) or the RNeasy mini kit (Qiagen, 74104), according to the manufacturer’s instructions. For gene expression analysis, 500-1000 ng of total RNA were treated using RQ1 RNase-Free DNase (Promega, M6101) and retro-transcribed using the Maxima H Minus First Strand cDNA Synthesis Kit (ThermoFisher Scientific, K1652). RT samples were diluted and used as the template DNA for RT-qPCR using the iTaq™ Universal SYBR® Green Supermix (Biorad, 1725124) on either the QuantStudio™ 5 Real-Time PCR System or ViiA 7 Real-Time PCR System real-time PCR system. List of used oligos in Table S9.

### Mass spectrometry data analysis

All raw data was analyzed using MaxQuant (version 2.5.2.0) as previously described.(78) Search was performed against an *in-silico* digested UniProt reference proteome for *Homo sapiens* including canonical and isoform sequences (29th August 2022). Database searches were performed according to standard settings with the following modifications: Digestion with Trypsin/P was used, allowing 2 missed cleavages. Oxidation (M), acetyl (protein N-term) allowed as variable modifications with a maximum number of 2. Carbamidomethyl (C) was disabled as fixed modification. Label-Free Quantification (LFQ) was enabled, not allowing Fast LFQ. Match between runs was enabled on default settings. Output from MaxQuant was exported and processed for statistical analysis in the Perseus computational platform version 1.6.14.0.(79) LFQ intensity values were log2 transformed and potential contaminants and proteins either identified by site or only reverse peptides were removed. Samples were grouped in experimental categories and proteins not identified in 4 out of 4 replicates in at least one group were removed. Missing values were imputed using normally distributed values with 0.3 width and 1.8 down shift for the total matrix. After imputation, statistical analysis was performed using two-sided Student’s *t* tests. Results were exported into in MS Excel 365 for a comprehensive browsing and visualization of the datasets. Volcano plots were constructed for data visualization using the VolcaNoseR web app (80).

### Metabolomics

MOLM-13 knock-down cell lines (shCONTROL and shBCLAF1) were induced for 96h with DOX in the usual RPMI culturing media. Then, cells were pelleted, washed, and seeded in RPMI no glucose (11879020) containing 11 mM ^13^C_6_-glucose (CLM-1396-2 Cambridge Isotope Laboratories) for 24 hours. After cells were quenched and metabolites were extracted as previously described.(81) In brief, cells were quickly pelleted by centrifugation (1 min at 1500 x g), followed by two quick washes (1 min at 1500 x g each) with -40 °C-cold quenching buffer (10 mM ammonium acetate in 60:40 methanol-water). The quenched cell pellets were then frozen on dry ice, and stored at - 80 °C until extraction. Metabolite extraction was performed using cold two-phase methanol–water–chloroform extraction containing 0.75 mg/mL glutaric acid (Sigma-Aldrich) as internal standard. After vortexing for 10 min, phase separation was achieved by centrifugation at 4 °C for 10 min, and methanol phase (containing the total polar metabolites content) was separated and dried by nitrogen evaporator. For metabolite detection and quantification, gas chromatography–mass spectrometry (GC-MS) analysis was performed as previously described. (82) Briefly, polar metabolite phase was derivatized in 20 µL of 20 mg/ml methoxyamine in pyridine per sample for 90 min at 37 °C. Subsequently, 15 µL of N-(tertbutyldimethylsilyl)-N-methyl-trifluoroacetamide, with 1 % tert-butyldimethylchlorosilane were added to 7.5 µL of each derivative and incubated for 60 min at 60 °C. The metabolites were separated by gas chromatography (Agilent 8860 GC system), with an inlet temperature set at 270 °C and a 1 µL spitless sample injection on a DB35MS column (30 m, 0.25mm, 0.25 µm). Elution was carried out using a constant 1 ml/min flow of helium as gas carrier and gradient of temperature. The column temperature was initially held at 100 °C for 1 min and increased with a 2.5 °C/min slope until reaching and holding 105 °C for 2 min, then further increased with a 3.5 °C/min slope until reaching 240 °C, and finally increased with a 3.5 °C/min slope until reaching and holding 320°C for 2 min. After eluting from the column, the polar metabolites were detected by mass spectrometry (Agilent 5977C MS system), using electron impact ionization with source and quadrupole temperatures set at 230 °C and 150 °C, respectively. The peak area was subsequently normalized to the protein content of the sample or total cell count and to the internal standard glutaric acid. MATLAB was used to extract mass distribution vectors and integrated raw ion chromatograms. The natural isotopes distributions were corrected considering a combination of the theoretical mass distribution vectors (MDVs) for all potentially labeled isotopologues of a metabolite. (83)

### RNA-seq

Inducible knockdown cell lines (shCONTROL and shBCLAF1) were treated with doxycycline for 96h. RNA was extracted using the Direct-zol RNA MiniPrep Kit (Zymo Research, R2050), according to the manufacturer’s instructions. Three independent biological replicates were collected for each condition. RNA libraries were prepared and sequenced at the CABIMER Genomics Facility (CABIMER, Sevilla). The used kit for library preparation was the Stranded Total RNA Preparation Ribo-Zero kit (Illumina), and sequencing was performed with the NovaSeq 6000 SP reagent kit (Illumina) with 2 × 50 bp paired-end reads in a NovaSeq 6000 system (Illumina).

### RIPseq and RIP-qPCR

Inducible knockdown cell lines were collected after 96 h of doxycycline treatment for BCLAF1-RIP-seq using anti-BCLAF1 antibody. Cell pellets were lysed using PEB buffer (20 mM Tris-HCl pH 7.5, 100 mM KCl, 5 mM MgCl_2_, 10% NP-40, Cocktail Complete (Sigma-Aldrich, 11697498001), PMSF (Millipore, 52332) and 5 µL RNasin® Ribonuclease Inhibitor (Promega, N2111) every 1 mL of buffer). The IP was set using 300 µL NT2 buffer (50 mM Tris-HCl pH 7.5, 150 nM NaCl, 1 mM MgCl_2_, 0.05% NP-40) complemented with 1 mM DTT, RNasin and proteinase inhibitors, 1.5 mM EDTA and protein extract up to 1 mL. For each IP, 2 µg of anti-BCLAF1 antibody were used and left overnight at 4°C on rotation. The primary antibody was previously attached to protein A/G dynabeads for 1 h at RT on rotation. Beads were washes with NT2 buffer before adding them to the protein lysate. DNase treatment (Invitrogen, AM1907) was done on beads for 10 min at 37°C. Afterwards, RNA elution from the beads and protein digestion was accomplished with a Proteinase K treatment (ThermoFisher, 25530049) for 20 min at 55°C on agitation. The eluted sample follows RNA purification using Trizol and following manufacturer’s instructions. RNA was then used either for library preparation for RIP-seq (as indicated in the RNA-seq segment) or retro-transcribed using Maxima H Minus First Strand cDNA Synthesis Kit (ThermoFisher Scientific, K1652) for RIP-qPCR analysis. 10% of the protein extract that was used for the IP was used as INPUT and extracted with Trizol.

### RNA-seq data analysis

Reads were aligned using STAR v2.7.11b (84) and mapped to hg38 reference human genome. Quantification of the reads was performed using StringTie v2.2.3.(85) Downstream analysis of the data was conducted with custom Rscripts (https://github.com/gmzlab/Chromatin_Modification_scripts), under R v4.3.2. Differential gene expression analysis was performed using DESeq2 v1.42.1 from Bioconductor. (86) Differentially expressed genes were determined by FC ≥ 2 and FDR < 0.05 for upregulated genes and FC ≤ -2 and FDR < 0.05 for downregulated genes. For RIP-seq data, BCLAF1-bound RNAs were determined using DESeq2 as well (FC ≥1.5 and FDR < 0.05). For further either gene ontology (GO) enrichment analysis or Gene Set Enrichment Analysis (GSEA), clusterProfiler v.4.10.1. (87) The threshold to determine significative gene ontology terms was FDR < 0.05. Every venn diagram was performed with R package VennDiagram v1.7.3.

### Publicly available data analysis

RNA-seq data from TCGA cohorts was downloaded using TCGAbiolinks v2.30.4. (88) Downstream analysis were performed using a custom Rscript which uses the same packages as the one used for RNA-seq (https://github.com/gmzlab/Chromatin_Modification_scripts). Differentially expressed genes and gene ontology terms were determined by the same thresholds fixed in our RNA-seq data. The cohorts used were: BEAT-AML (28) and TARGET-AML.(89) ENCODE ChIP-seq datasets for ATF4 in K562 cells were downloaded from the GEO (GSE127432). Reads were mapped using BWA v0.7.17,(90) downstream analysis samtools v1.13 and peak calling using MACS3 v3.0.2. (91)

### Statistical analysis

All data in bar- and line-plots are presented as mean ± SEM of at least three independent replicates, unless stated otherwise. Statistical significance was analyzed using two-way analysis of variance ANOVA (two-way ANOVA), unpaired or paired *t* test at a confidence interval of 95%, unless otherwise specified. All statistical analyses were performed using GraphPad Prism software. \**p* < 0.05, \*\**p* < 0.01 and \*\*\**p* < 0.001 were considered significant.

## Data sharing

Proteomics data are available at PRIDE (PXD065945). RNA-seq and RIP-seq data are available at GEO (GSE303836).

To review PRIDE accession PXD065945, log in to the PRIDE website using: Project accession: PXD065945

Token: rfnN7KFJkLhS

To review GEO accession GSE303836: o to https://www.ncbi.nlm.nih.gov/geo/query/acc.cgi?acc=GSE303836 Enter token slwrqyacpponrsp into the box

## Acknowledgments

We thank the Genomic, Flow Cytometry and Microscopy Core facilities at CABIMER for their technical help, and A.J. Bannister for critically reading the manuscript. G.M.-Z. was supported by a fellowship from “la Caixa” Foundation (ID 100010434) and from the European Union’s Horizon 2020 research and innovation programme under Marie Skłodowska-Curie grant agreement No. 847648. The fellowship code is LCF/BQ/PR21/11840007. The G.M.-Z. laboratory was supported by grants PID2021-127432NA-I00 and PID2023-151942NB-I00 funded by MCIN/AEI/10.13039/501100011033 and by FEDER, UE, and by grant CNS2022-135600 funded by MICIU/AEI/10.13039/501100011033 and by European Union NextGenerationEU/PRTR. This work was supported by the National Institutes of Health (NIH) grants R01AI173077 (J.J.B.), T32 AI007163 (S.J.C.), F31HL165908-01A1 (S.J.C.), and R01HL152180 (J.A.M.). J.J.B. was supported by the American Society of Hematology, St. Louis Children’s Hospital Foundation, the Children’s Discovery Institute, Hyundai Hope on Wheels, and Kelsie’s Hope Foundation. This publication was supported by the Washington University NIH/NCI SPORE in Leukemia Grant 1P50CA171063 (J.J.B.). J.A.M. is a Scholar of the Leukemia and Lymphoma Society. The R.G.-P. laboratory was supported by the EMERGIA 2020 program (EMERGIA20_00276) from the Consejería de Economía, Conocimiento, Empresas y Universidad, Junta de Andalucía. L.P. and S.G. were supported by intramural IIT funding. The I.B. laboratory was supported by the AIRC Startup grant 26505. This manuscript is the result of funding in part by the NIH. It is subject to the NIH Public Access Policy. Through acceptance of this federal funding, NIH has been given a right to make this manuscript publicly available in PubMed Central upon the Official Date of Publication, as defined by NIH. This publication is solely the responsibility of the authors and does not necessarily represent the official view of NIAID, NCI, NCRR, or NIH.

## Authorship Contributions

Conceptualization: J.J.B and G.M.-Z.; Formal analysis: L.L.-H., R.A. and L.P.; Funding acquisition: S.G., R.G.-P., I.B., P.A.-M, S.J.C., J.J.B., and G.M.-Z.; Investigation: L.L.-H., N.K., S.C.-S., S.J.C., P.T.-K., D.I., L.W., P.A.-M, and E.S.-H.; Consultation: J.A.M.; Project administration: R.G.-P., L.P., I.B., P.A.-M, J.J.B., and G.M.-Z.; Software: L.L.-H., R.A. and L.P.; Supervision: R.G.-P., L.P., I.B., J.J.B., and G.M.-Z.; Visualization: L.L.-H. and S.J.C. Writing – original draft: G.M.-Z. Writing – review & editing: L.P., I.B., L.L.-H., S.J.C., J.J.B., and G.M.-Z.

