## Supplemental Figures for "BCLAF1 links RNA splicing to ATF4-dependent metabolic adaptation in acute myeloid leukemia"

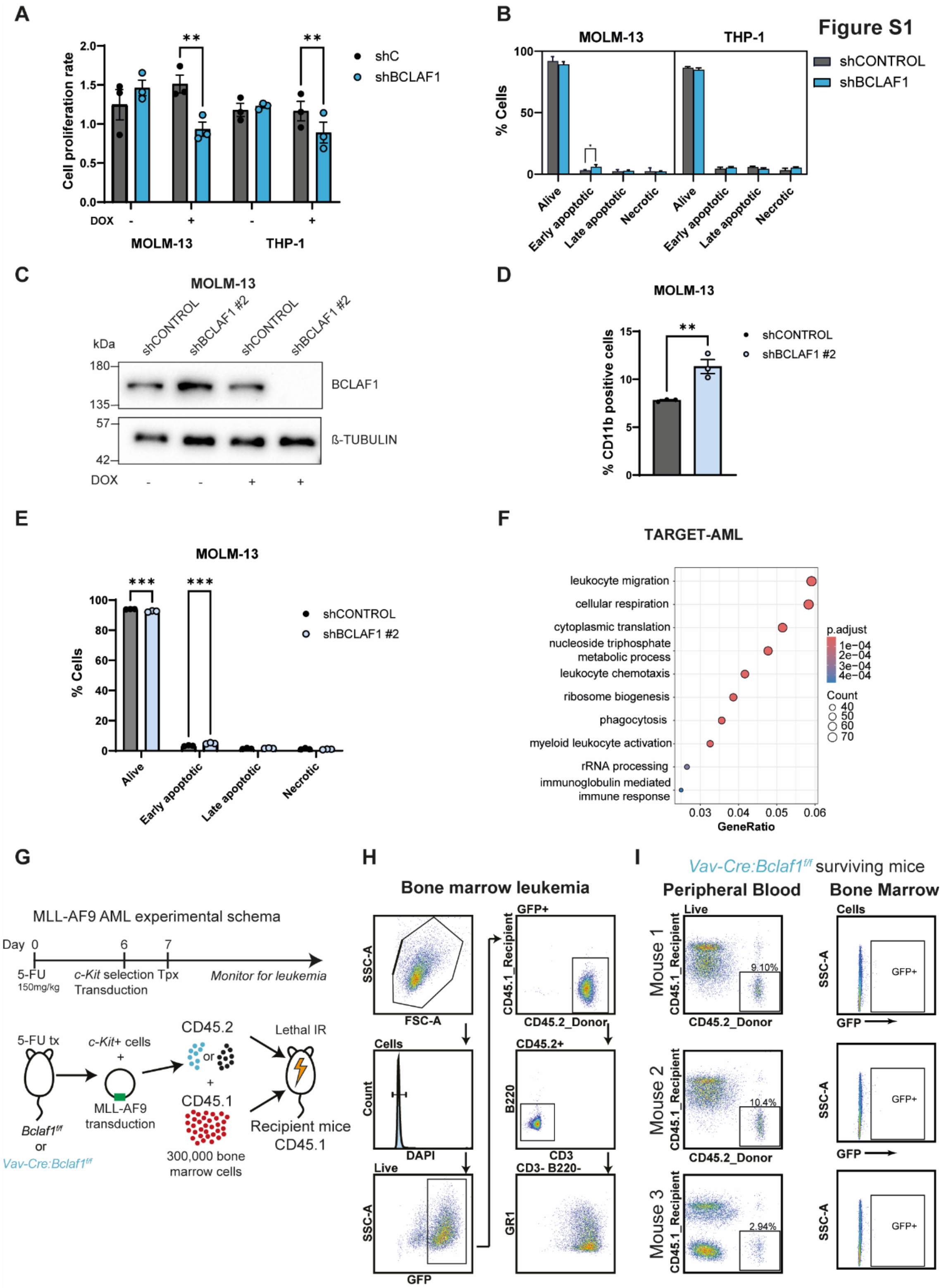


**Figure S1. Supplementary to BCLAF1 supports AML pathogenesis. A.** Cell proliferation assay (MTT-based) in shCONTROL and shBCLAF1 cells treated or not with DOX for 96 hours, followed by 96 additional hours in culture. Data represent mean ± SEM of three biological replicates (**p < 0.01; paired t-test). **B.** Annexin V/PI flow cytometry analysis showing the distribution of live, early apoptotic, late apoptotic, and necrotic cells following 96h of DOX treatment. Data represent mean ± SEM of three biological replicates (*$p < 0.05$; two-way ANOVA). **C.** Immunoblot analysis of BCLAF1 protein levels in shCONTROL and shBCLAF1 #2 cells treated or not with DOX for 96 hours. β-tubulin was used as a loading control. **D.** Flow cytometry analysis showing the percentage of CD11b-positive cells in shCONTROL and shBCLAF1 #2 cells treated with DOX for 96h. Data represent mean ± SEM of three biological replicates (**$p < 0.01$; unpaired t-test). **E.** Annexin V/PI flow cytometry analysis showing the distribution of live, early apoptotic, late apoptotic, and necrotic cells following 96h of DOX treatment. Data represent mean ± SEM of three biological replicates (***$p < 0.001$; two-way ANOVA). **F.** Dot plot of GO terms enriched among genes up-regulated in BCLAF1-low versus BCLAF1-high patient samples from the TARGET-AML cohort. GeneRatio indicates the proportion of significantly upregulated genes associated with each GO term. Dot size reflects the number of genes per term; color denotes adjusted p-value. **G.** Schematic of in vivo leukemia experiments in Figure 1G using mice with hematopoietic-specific loss of *Bclaf1*. HSPCs from CD45.2+ *Bclaf1f/f* or *Vav-Cre:Bclaf1f/f* mice were transduced with retrovirus co-expressing MLL-AF9 and GFP then transplanted with 300,000 CD45.1+ recipient bone marrow cells into lethally-irradiated CD45.1+ recipient mice. Leukemic burden was assessed by percentage of GFP+ cells in peripheral blood. Mice were euthanized once peripheral blood showed ≥ 80% leukemia. **H.** Representative flow cytometry gating for bone marrow of mice with leukemia from Figure 1G. Leukemia cells were defined as GFP+. **I.** Flow cytometry of peripheral blood and bone marrow from representative surviving mice that received CD45.2+ *Vav-Cre:Bclaf1f/f* HSPCs transduced with MLL-AFP- and GFP-expressing retrovirus. Peripheral blood shows percentage of CD45.2+ donor cells in all live cells. Bone marrow shows percentage of GFP+ leukemia cells among all leukocytes. Data are from 3 independent mice that were alive at 154 days post-transplant in Figure 1G.

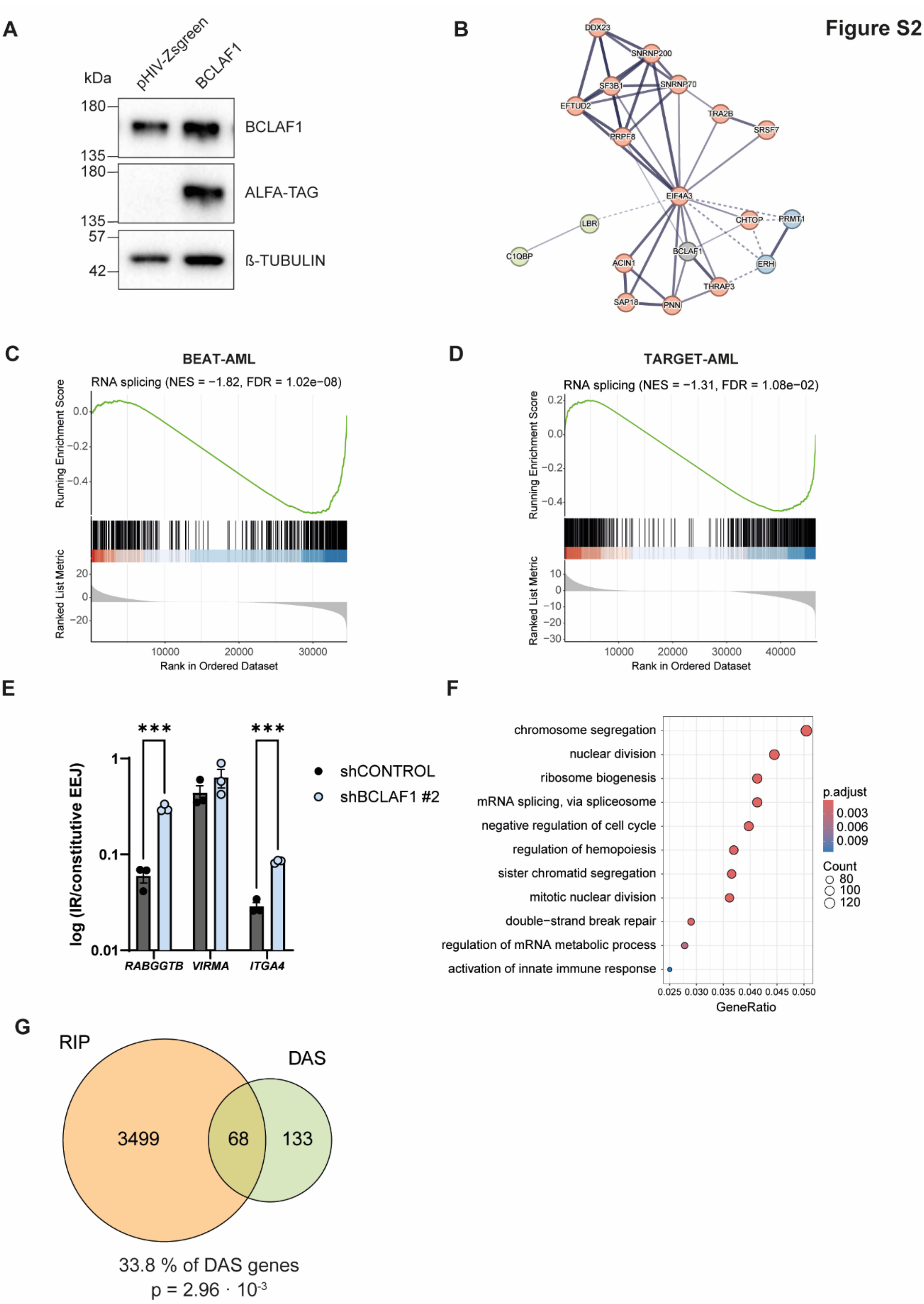


**Figure S2. Supplementary to BCLAF1 regulates mRNA splicing in AML cells. A.** Immunoblot analysis of MOLM-13 cells infected with a lentiviral vector expressing ALFA-tagged BCLAF1 or empty vector control (pHIV-Zsgreen). Blots were probed with anti-BCLAF1 and anti-ALFA antibodies. β-tubulin was used as a loading control. **B**. STRING network analysis of BCLAF1 interactors (n = 19) identified in panel A. Grey circle indicates BCLAF1; red circles represent spliceosome components. **C.** and **D.** Gene set enrichment analysis (GSEA) of RNA splicing–related gene sets in BCLAF1-low versus BCLAF1-high AML patient samples from the BEAT-AML and TARGET-AML cohorts, respectively. **E.** RT-qPCR experiments showing relative intron retention (IR) levels of the indicated transcripts in shCONTROL and shBCLAF1 #2 cells treated with DOX for 96 hours. Data represent mean ± SEM of three biological replicates (***$p < 0.001$; unpaired t-test). **F.** Dot plot of GO terms enriched among BCLAF1 interactors. Dot size reflects the number of genes per term; color denotes adjusted p-value. **G.** Venn diagram showing the overlap between BCLAF1-bound transcripts (RIP), transcripts with differential alternative splicing (DAS) upon BCLAF1 knockdown, and genes downregulated upon BCLAF1 depletion (RNA-seq). Only genes annotated in amino acid metabolism-related GO categories were considered among the downregulated set.

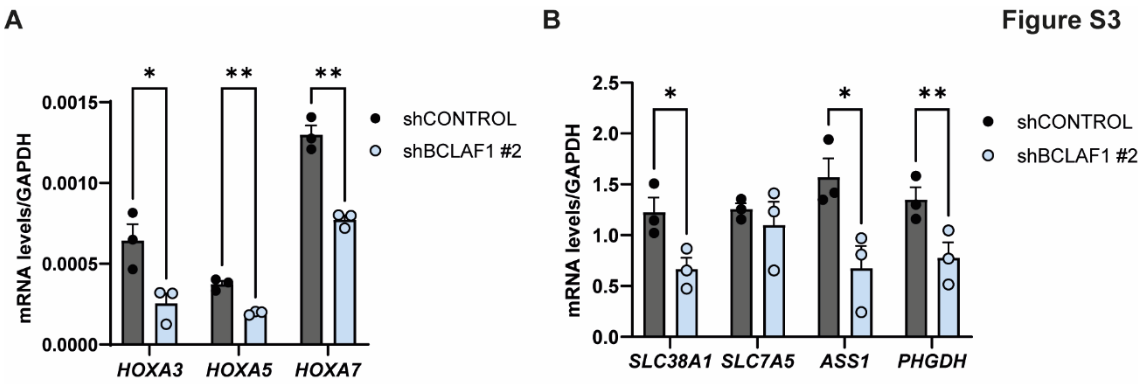


**Figure S3. Supplementary to BCLAF1 promotes amino acid metabolic gene expression. A.** and **B.** RT-qPCR experiments showing relative mRNA levels of the indicated transcripts in shCONTROL and shBCLAF1 #2 cells treated with DOX for 96 hours. Data were normalized to *GAPDH* and represent mean ± SEM of three biological replicates (*p < 0.05, **p < 0.01; unpaired t-test).

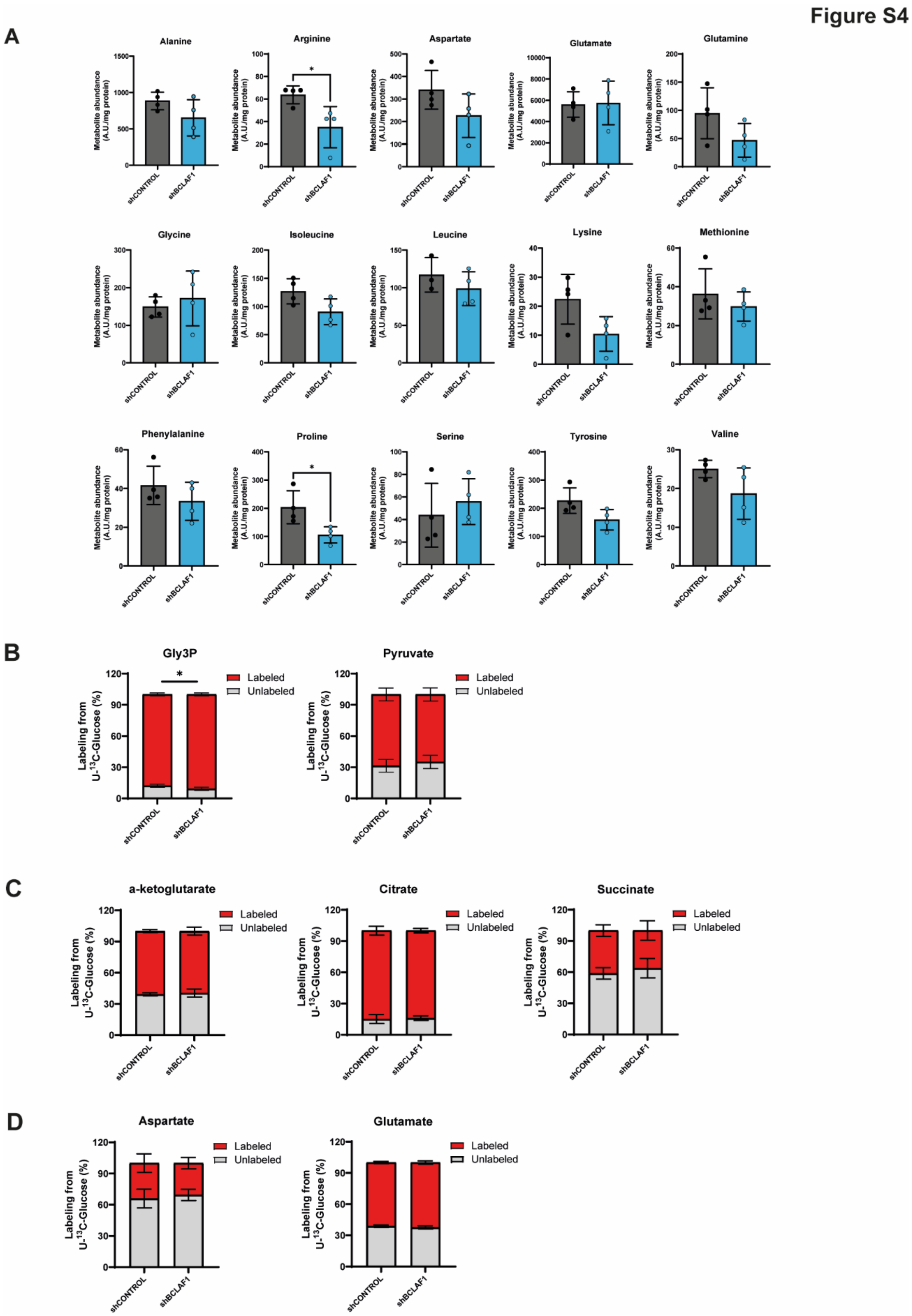


**Figure S4. Supplementary to loss of BCLAF1 disrupts *de novo* amino acid synthesis. A.** Barplots showing total amino acid levels in shCONTROL and shBCLAF1 cells treated with DOX for 96 hours. Data are shown as mean ± SEM of four technical replicates from one representative experiment (*$p < 0.05$; unpaired t-test). The experiment was independently repeated with similar results. **B-D.** Percentage of unlabeled (M+0; gray) or labeled (the sum of M+1, M+2, and M+3; red) metabolite levels. Data are shown as mean ± SEM of four technical replicates from one representative experiment (*$p < 0.05$; two-way ANOVA). The experiment was independently repeated with similar results.

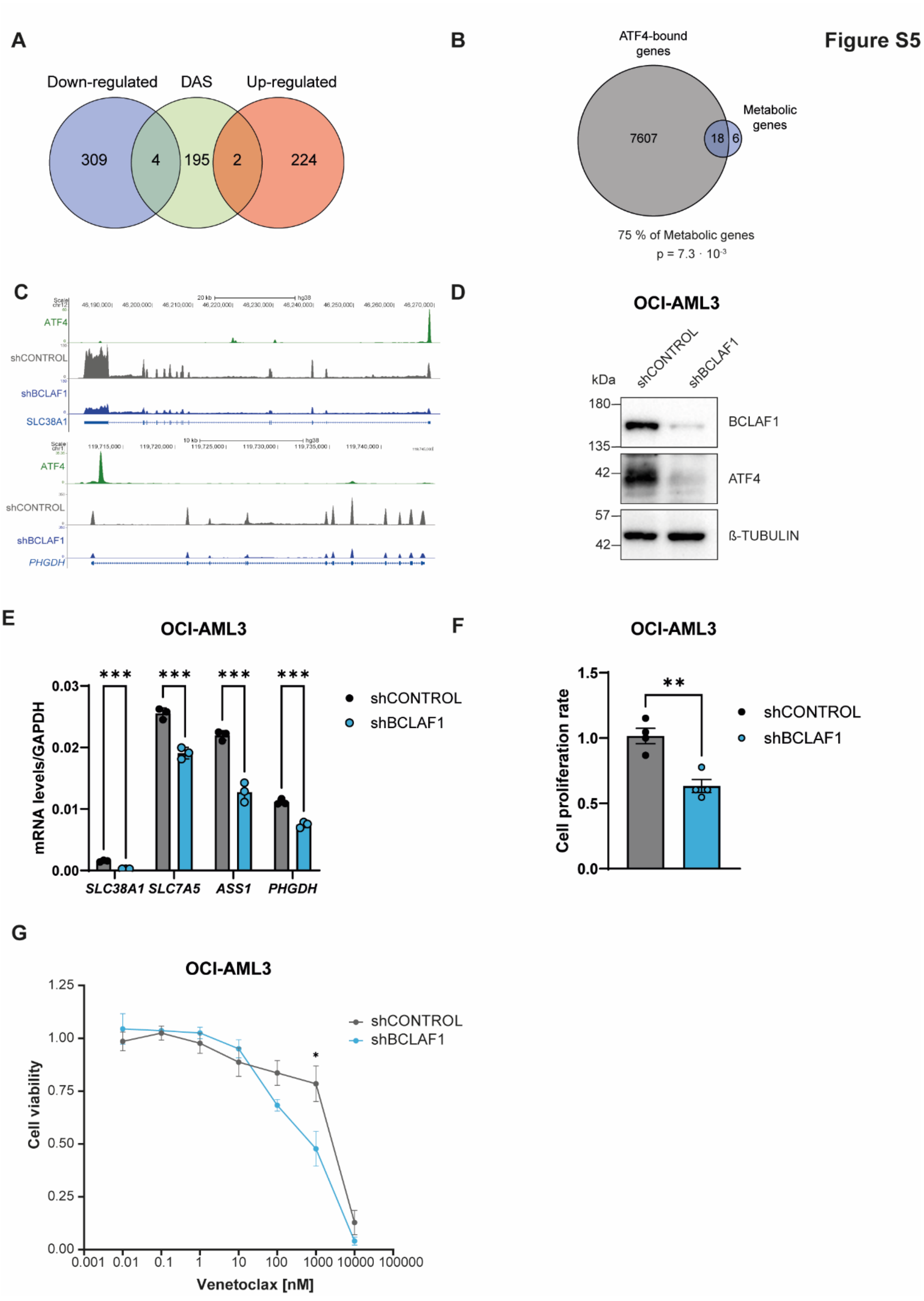


**Figure S5. Supplementary to BCLAF1 supports amino acid metabolism through ATF4. A.** Venn diagram showing the overlap between up-regulated, down-regulated and differentially spliced genes (DAS) upon BCLAF1 depletion. **B.** Venn diagram showing the overlap between genes downregulated upon BCLAF1 depletion (RNA-seq) and genes with significant ATF4 ChIP-seq peaks (ENCODE, K562 cells). Only genes annotated in amino acid metabolism-related GO categories were considered among the downregulated set. Overlap significance was assessed by Fisher's exact test. **C.** Genome browser snapshots showing ATF4 ChIP-seq signal from ENCODE data in K562 cells together with RNA-seq data from shCONTROL and shBCLAF1 cells treated with DOX for 96 hours. **D.** Immunoblot analysis of BCLAF1 and ATF4 protein levels in shCONTROL and shBCLAF1 cells treated or not with DOX for 96 hours in OCI-AML3 cells. β-tubulin was used as a loading control. **E.** RT-qPCR experiments showing relative mRNA levels of the indicated transcripts in shCONTROL and shBCLAF1 cells treated with DOX for 96 hours. Data were normalized to *GAPDH* and represent mean ± SEM of three biological replicates (***p < 0.001; unpaired t-test). **F.** Cell proliferation assay (CellTiter-Blue-based) in shCONTROL and shBCLAF1 cells treated with DOX for 96 hours, followed by 96 additional hours in culture. Data represent mean ± SEM of three biological replicates (**p < 0.01; paired t-test). **G.** Cell viability assay (CellTiter-Blue-based) in shCONTROL and shBCLAF1 cells treated or not with DOX for 96 hours, followed by 96 additional hours of Venetoclax treatment. Data is normalized to control (DMSO) and represent mean ± SEM of four biological replicates (*p < 0.05; unpaired t-test).

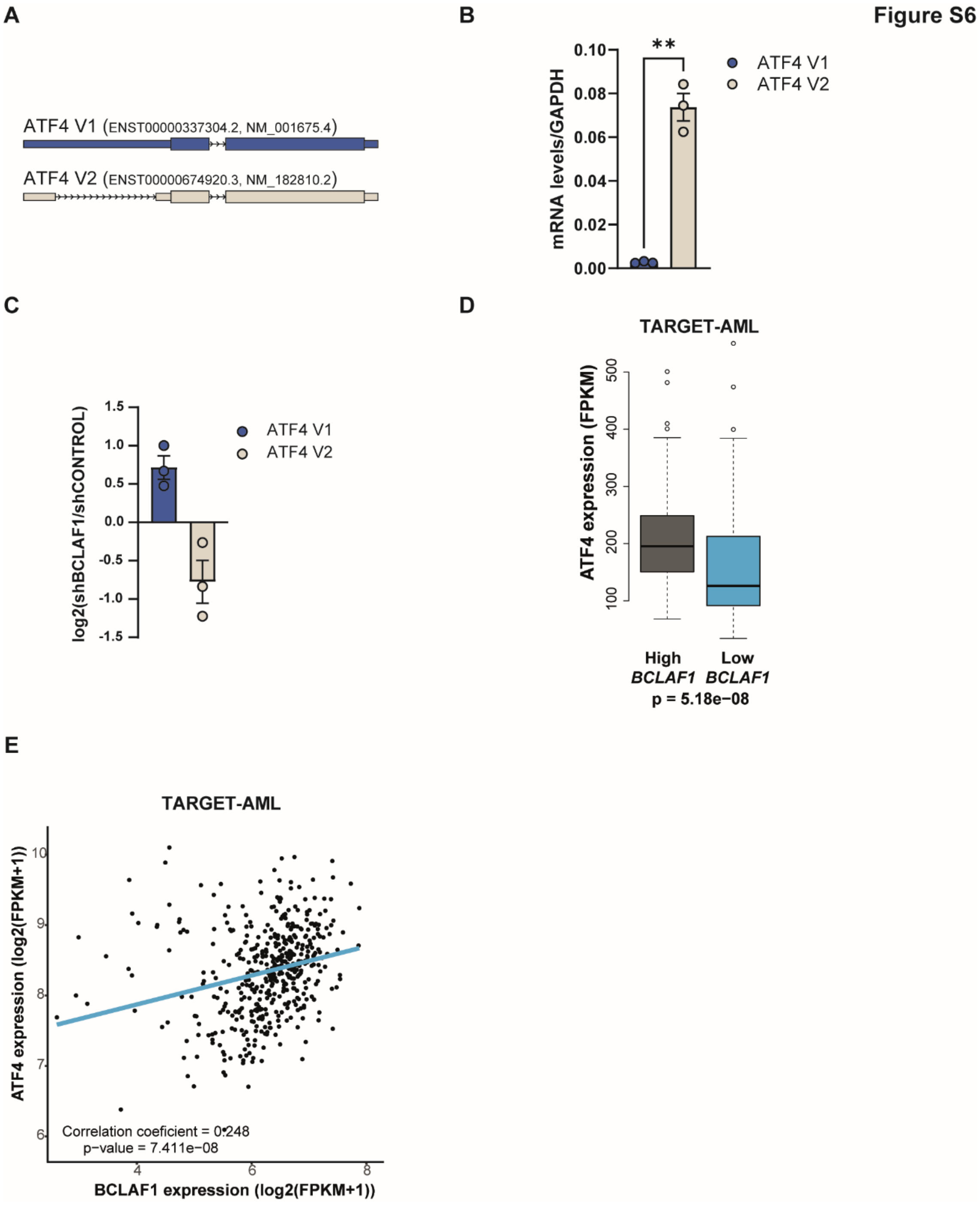


**Figure S6. Supplementary to BCLAF1 controls *ATF4* mRNA alternative splicing. A.** Schematic representation of both ATF4 isoforms being V1 the one that has an intron retention and V2 the functional one. **B.** RT-qPCR experiments showing relative mRNA levels of the indicated transcripts in shCONTROL cells treated with DOX for 96 hours. Data were normalized to *GAPDH* and represent mean ± SEM of three biological replicates (**p < 0.01; unpaired t-test). **C.** RT-qPCR experiments showing the ratio of relative mRNA levels of the indicated transcripts between shBCLAF1 and shCONTROL cells treated with DOX for 96 hours. Data were normalized to *GAPDH* and represent+ log2 of the ratio from both samples and ± SEM. **D.** Box plot showing ATF4 mRNA levels in BCLAF1-high (top 25%) and BCLAF1-low (bottom 25%) patient samples from the TARGET-AML cohort. Statistical significance was assessed using Wilcoxon Rank Sum and Signed Rank Tests (one-sided). **E.** Scatter plot showing the correlation between BCLAF1 and ATF4 mRNA levels across TARGET-AML patient samples. Pearson correlation coefficient (r) and p-value are indicated.
